# Phenotypic variability between closely related *Trichodesmium* strains as a means to making the most out of heterogeneous microenvironments

**DOI:** 10.1101/2024.05.22.595369

**Authors:** Mina Bizic, Danny Ionescu, Siyuan Wang, Futing Zhang, Yeala Shaked

**Affiliations:** The Leibniz Institute of Freshwater Ecology and Inland Fisheries, Dep. Plankton and Microbial Ecology, Stechlin, Germany; Technical University of Berlin, Dep. Environmental Microbiomics, Berlin, Germnay; The Fredy and Nadine Herrmann Institute of Earth Sciences, Hebrew University of Jerusalem, Jerusalem, Israel; The Interuniversity Institute for Marine Sciences in Eilat, Eilat, Israel

## Abstract

The cynobacteria *Trichodesmium* spp., play an important role in oceanic carbon and nitrogen cycles. Forming colonies that host diverse microorganisms likely enhances Trichodesmium’s metabolic flexibility and resilience. Combining single-colony metagenomics and metatranscriptomics with a novel bioinformatic approach we investigated the community composition and transcriptional activity of 69 individual *Trichodesmium thiebautii* colonies collected from the northern Red Sea, Israel. Three *Trichodesmium* strains were detected in all colonies, each with a heterogenous yet distinct transcriptional-activity profile. N-fixation, superoxide dismutase, and ATPase activities were similar among colonies, while P and Fe uptake, as well as photosynthesis, were high in only few colonies. Colonies had no core microbiome with >70% dissimilarity in the associated microbial communities, yet, the abundance of some species was correlated with the activity of specific *Trichodesmium* strains. We propose *Trichodesmium* occupies heterogeneous microenvironments with which it deals through inter-strain phenotypic variability further fine-tuned through interactions with beneficial, non-fixed, heterotrophic colonizers.

## Introduction

Trichodesmium spp. are filamentous colony-forming marine cyanobacteria of global significance ^1,2^. Several species of Trichodesmium are known, with noticeable genetic and functional differences between them ^3,4^. Thanks to its ability to fix nitrogen, in addition to CO_2_, *Trichodesmium* forms large surface blooms in tropical and sub-tropical waters ^5^. Hundreds of *Trichodesmium* filaments often aggregate as a colony, a form that offers several advantages^6^. *Trichodesmium* colonies are colonized by different microorganisms including heterotrophic bacteria, algae, fungi, and protists ^7–9^. These microorganisms have been suggested to aid the colony in acquiring iron from dust particles by producing iron-solubilizing molecules (siderophores), in exchange for photosynthetic exudates and leaking fixed nitrogen ^7,10^. It was further suggested that the assembly into colonies leads to reduced grazing ^11^.

Similar to other organic matter particles in aquatic environments, *Trichodesmium* colonies are often studied as bulk concentrates containing hundreds or thousands of colonies ^9,12,13^. Such analyses have pinpointed, for example, potential key interactions between the cyanobacteria and colonizing microorganisms ^14^, or different activity between large water masses^15^. Nevertheless, the study of individual OM particles in marine environments is a critical aspect of marine science, shedding light on the composition, sources, and fate of particulate organic matter (POM) in the ocean. Analyzing individual OM particles provides insights into the dynamics of carbon and nutrient cycling in marine ecosystems, and the succession of microorganisms responsible for these processes ^16^. It further allows deconvoluting the different sources and history of OM particles which lead to biased results ^17,18^.

Recent studies explored the variability among individual OM particles revealing a large heterogeneity in colonizing microorganisms and their activities ^17,19,20^. This heterogeneity was attributed to stochastic colonization of the particle, competitive interactions between colonizers, as well as variability in particle age and source. Accordingly, it emerges that individual particles in aquatic environments are each an individual micro-ecosystem ^18^. Similar variability was observed also in the photosynthetic activity of individual *Trichodesmium* colonies collected at the same time and place ^21,22^, as well as in dust particles processing ^23^.

We employed a novel approach combining metagenomics and metatranscriptomics of individual colonies to explore the variability in the composition and transcriptional activity of *Trichodesmium* colonies during a late autumn bloom in the Gulf of Eilat (Aqaba), the Red Sea, Israel. By studying individual colonies, we investigated the frequency of occurrence of different microorganisms within the colonies, and the correlation of these organisms with *Trichodesmium strain* diversity per colony. To better understand the links between *Trichodesmium* colonies and their surrounding environment, we compared the transcriptional activity of the individual colonies. We consider heterogeneity in transcriptional activity to reflect the micro-environment characteristics of a colony. In contrast, depending on the function, homogeneity in transcriptional activity points to an overarching property of the environment beyond the colony, or to a general function, essential for the survival of the cells.

## Results

### Community composition of individual *Trichodesmium* colonies

Out of the 100 individual *Trichodesmium* colonies collected over 5 consecutive days, we were able to amplify and sequence 69 colonies. The microbial communities colonizing the individual *Trichodesmium* colonies were heterogeneous with very low similarity between them regardless of the sampling day (Fig. 1). The majority of the data was bacterial, with a few colonies displaying a high abundance of viral reads (Fig. 1A). *Trichodesmium* was the most dominant species (Fig. 1B), yet the colonies consisted of several cyanobacterial species with *Okeania* being the most common after *Trichodesmium* (Fig. 1B). While some genomes previously annotated as *Okeania* were recognized as *Trichodesmium*, the annotated contigs matched *Okeania* genomes with <85 % average nucleotide identity to known *Trichodesmium* genomes and did not align to the Trichodesmium strains in the colonies (Table S1). The colonies were colonized by diverse *Bacteria* and *Archaea* (Fig. 1C-D), the most common of which belonging to the genera *Pelagibacter, Glivibacter*, and an unnamed *Pirellulaes* genus (UBA1268) (Fig. 1C). Most of the archaeal community belongs to the *Poseidoniaceae* family (Fig. 1D). Reads of eukaryotic microorganisms, present in lower abundance than *Bacteria* and *Archaea* (Fig. 1A), were dominated on most colonies by fungi and Oomycota. Overall, the similarity in community composition was very low between most colonies with no clustering based on the sampling day (Fig. 1F).

**Figure 1.**
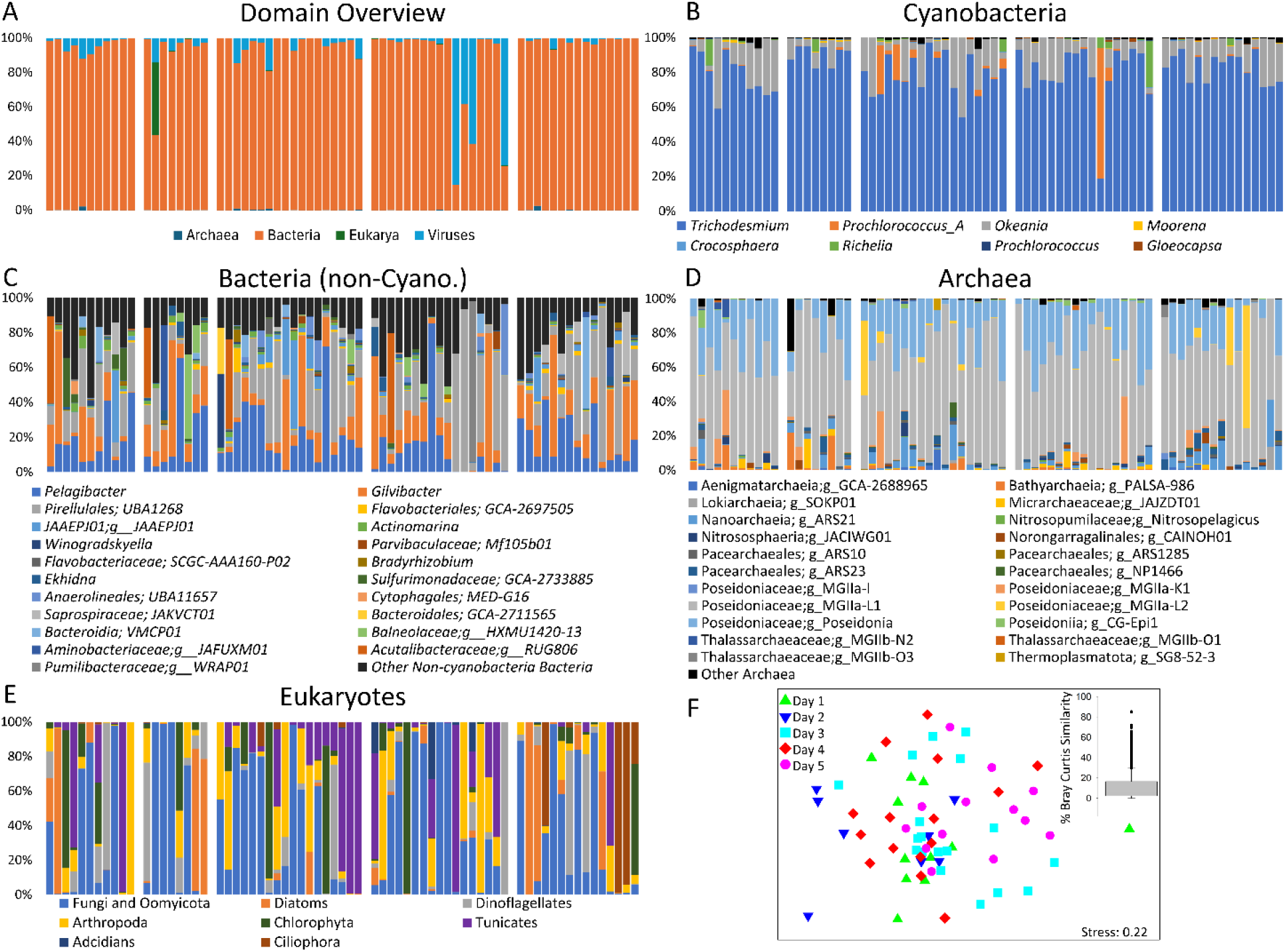
Microbial community composition on each of the *Trichodesmium* colonies. General abundance of *Bacteria, Archaea*, eukaryotes, and viruses as calculated from RAT analysis (A). The relative abundance of different cyanobacterial taxa, non-cyanobacteria *Bacteria*, and *Archaea* are shown in panels B-D, respectively as obtained from CAT analysis using the GTDB database for annotation. The relative abundance of eukaryotes results from RAT analysis using the NR database for annotation and is shown in panel (E). The heterogeneity in the microbial community found on *Trichodesmium* colonies is evident in a non-metric multidimensional scaling ordination (F) and in the bar charts (panels C-E). The insert in panel (E) depicts the distribution of similarities between the communities on different *Trichodesmium* colonies. Each bar in panels A-E represents an individual colony and is grouped according to sampling days.

Network analysis was conducted on the contig-based community analysis since 92-99 % of the raw data could be mapped to the assembled contigs (Fig. 2). This analysis revealed two main separate interaction networks with overall weak positive and negative interactions (Fig. 2). The first included *Bacteria* and *Archaea* associated with *Trichodesmium, Okeania*, and other *Cyanobacteria*, with no apparent interaction between the non-cyanobacterial species among themselves. The other network did not include any *Cyanobacteria* and revealed multiple interactions between the organisms, with the most connected nodes belonging to the genus *Hyphobacterium*, and the unnamed *Flavobacteriales* genus, GCA_002715885. No strong associations were observed between colony-associated *Bacteria* or *Archaea* and *Trichodesmium*, or between the associated prokaryotes themselves.

**Figure 2.**
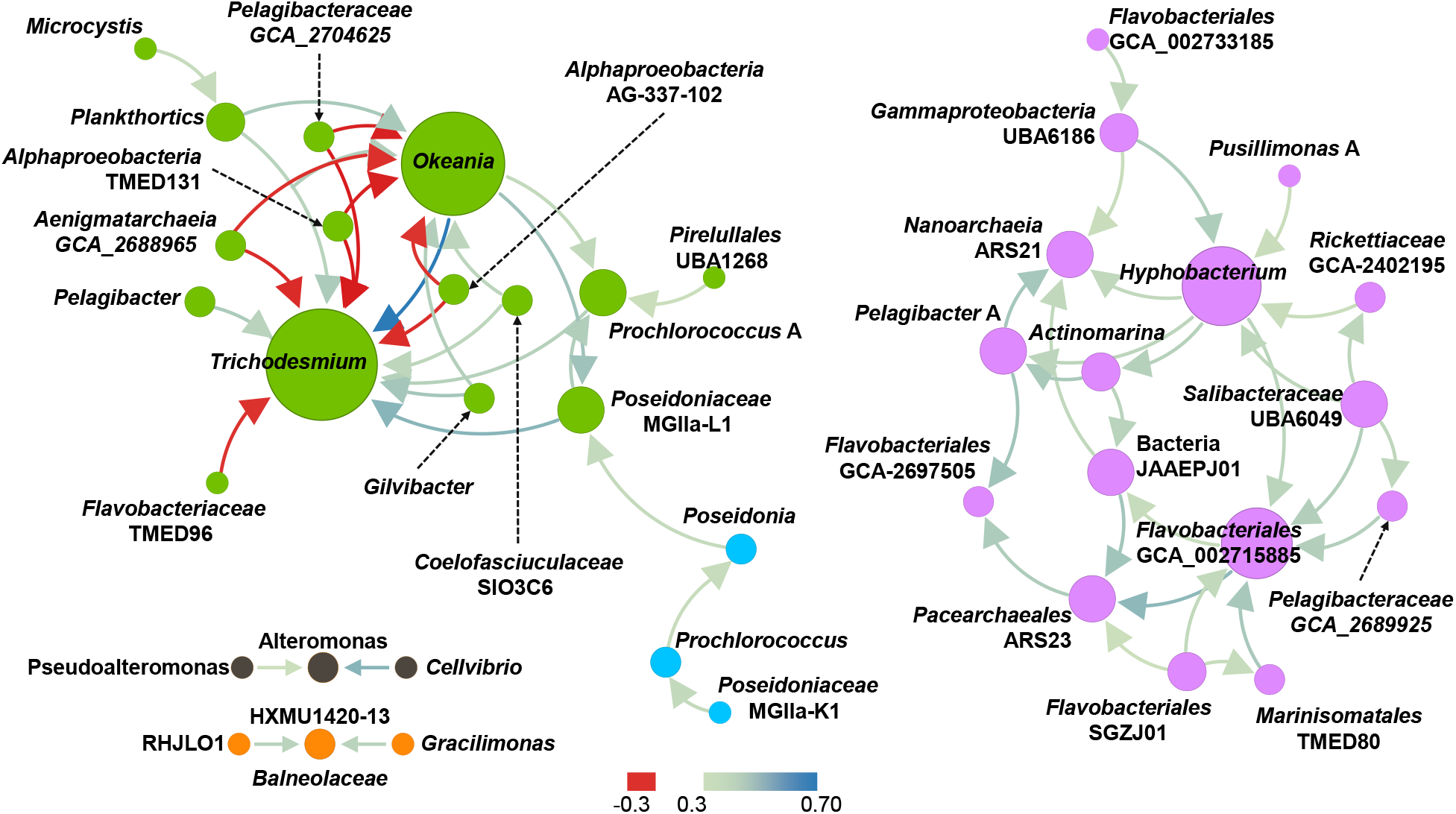
Network analysis of prokaryotic taxa at the genus level using taxonomic annotation of contigs based on the GTDB database and the CAT tool. Edges (arrows) are colored on a gradient between red (negative association) and blue (positive association). Node size and color represent the degree of connectivity, and modularity, respectively. No strong associations were detected between *Trichodesmium* and any colony-associated *Bacteria* or *Archaea*.

### *Trichodesmium* diversity

We obtained multiple metagenome-assembled genomes of *Trichodesmium* sp. Upon dereplication, we retained 3 MAGs with completeness values above 89 % and an average nucleotide identity of 99 %, representing strains of *Trichodesmium thiebautii*, (Fig 3A). Quantification of the relative abundance of the 3 *Trichodesmium strains* as well as *T. erythraeum* (IMS101) which is known from the region, revealed a variable distribution in each of the 69 colonies with absolute dominance of *T. thiebautii* strains and a low abundance of *T. erythraeum (*Fig. 3b). *T. miru*, also identified in this region ^12^, was not found in our data. Among the 3 strains of *T. thiebautii*, strain TRA20 was dominant in almost all colonies; however, strains TRA10 and TRC12 dominated some of the colonies as well (Fig. 3). In-Strain analysis ^24^ revealed further diversity within each of the three strains (Fig. S1).

**Figure 3.**
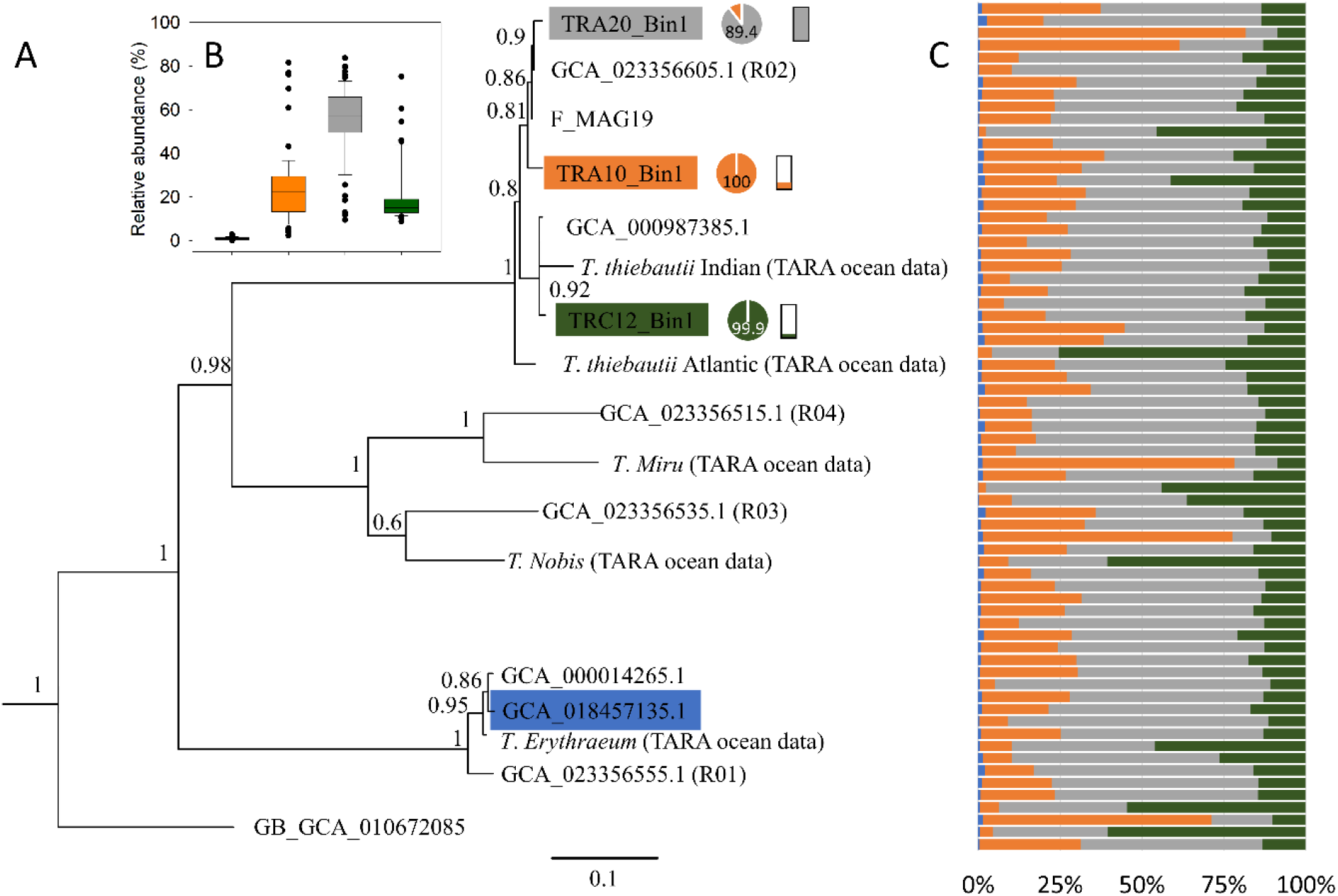
Maximum likelihood phylogenetic tree of *Trichodesmium* genomes based on multi-locus alignment of 120 marker genes as obtained from the GTDB-TK tool ^25^ (A). The distribution of the 3 different detected *Trichodesmium thiebautii* strains and *T. erythraeum* is summed up in panel (B) and detailed in panel (C). Completeness and contamination of the *Trichodesmium* metagenome-assembled genomes are shown in the pie and bar charts next to each genome in the tree. The full bar chart represents 7 % contamination (strain TRA20) whereas the lowest 0.93 % (strain TRC12).

Correlation analysis between the relative abundance of the 4 *Trichodesmium* sp. strains and that of bacterial and eukaryotic taxa present in at least 10 colonies revealed that only a few non-cyanobacterial taxa correlate with the abundance of different *Trichodesmium* strains (after FDR adjustment; Table S2), matching the results of the network analysis. Six genera were correlated with the abundance of strain TRC12, two with that of TRA10, and no taxon was correlated with the abundance of strain TRA20. *Gilvibacter* had the strongest correlation with the abundance of strain TRA10 (R=0.58 P_adj_=4e-5). Strain TRC12 was most strongly correlated with the abundance of two different strains of *Bacteroidia*, one belonging to the family *Crocinitomicaceae* (R=0.63, P_adj_=4e-6) and one to the unnamed family UBA9320 (R=0.59, P_adj_=2e-5). No taxon correlated with both *Trichodesmium* strains.

### Trichodesmium activity

Variability in the activity of *Trichodesmium* strains among the 69 individual colonies was tested on 195 (164 unique) selected genes belonging to key metabolic functions. These include P and Fe uptake, N fixation, photosynthesis carbon fixation, TCA cycle, circadian rhythm, ATP synthesis, cytochromes, pigment synthesis, and caspases. Some of the genes were not detected in all the strains while some were detected in multiple copies. The analysis revealed that the expression of some genes is highly variable among the colonies, while that of others is more homogenous (Fig. 4). This was confirmed using the Jensen-Shannon metric, comparing the expression pattern of each gene to a hypothetical homogenous expression across all colonies (Fig. 4), and with Shannon evenness (Fig. 5a). Grouping the genes into functional categories (Fig 5), revealed that general functions such oxygen radical detoxification (Superoxide dismutase; SOD), N fixation, the TCA cycle, and the circadian clock genes, are significantly more homogenous, for example, than nutrients uptake, photosynthesis, and pigment synthesis (Fig. 5). The latter result was obtained using both evenness (Fig. 5A) and the Jensen-Shannon metric (Fig. 5B), methods which rely on different statistical approaches to the data. Nevertheless, a large variation was observed between the expression patterns of genes included among the functions displaying a larger heterogeneity.

**Figure 4.**
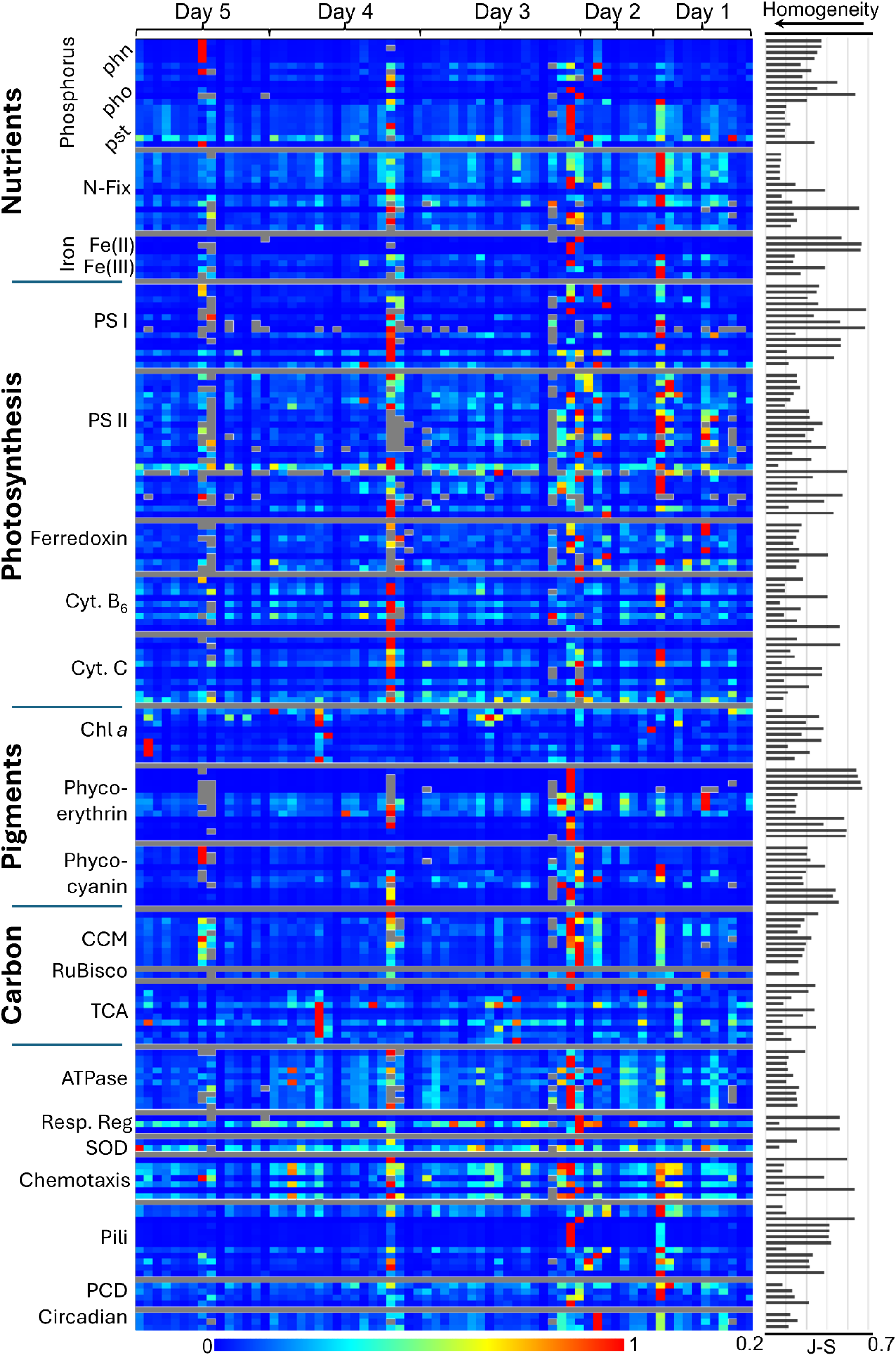
Gene expression of strain TRC12 for each of the 69 colonies. The gene expression is derived from the TPM (Transcript per kilobase million) counts for each gene normalized to the TPM counts of *rpo*B. The expression values for each gene were normalized such that the highest expression value equals 1. The Jensen-Shannon metric is provided for each gene and was derived from comparing the expression distribution for each gene and a homogenous vector of 69 data points each equaling 1. The lower the value, the more homogenous the gene expression is across the colonies.

**Figure 5.**
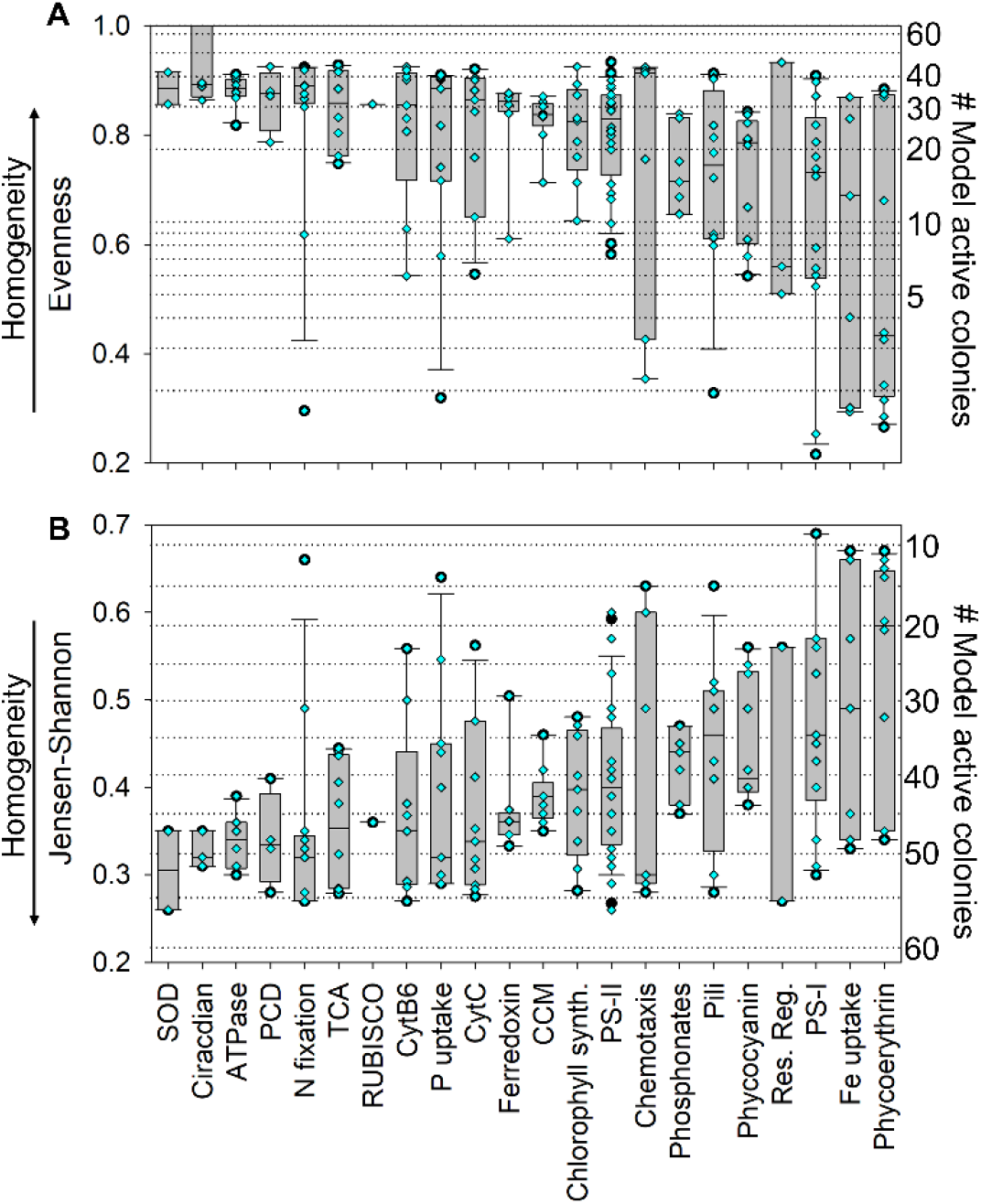
Heterogeneity of gene expression in strain TRC12 across deferent functional categories using Shannon evenness (A) or the Jensen-Shannon metric (B). The higher the evenness or lower the Jensen-Shannon value, the more homogenous the expression of the genes across the 69 colonies. For comparison, a scale was added on the right-hand side Y axis. The scale was generated using an increasing number of colonies (values on the axis) with an equal activity of 1. The rest of the colonies (up to 69) were set to an activity of 0.001. A different number of colonies was used for each index due to the behavior of the function. The arrow parallel to the left Y axis indicated the direction of increasing homogeneity.

A confirmation on the validity of the rpoB-normalization approach was obtained through the analysis of a public *Trichodesmium* transcriptome. The dataset from Cerdan-Garcia et al ^15^ (provided by the authors) was normalized to the RPKM values of the *rpo*B genes resulting in a similar clustering of samples and genes as obtained in their paper using the raw data set (Fig. S2).

We investigated the co-expression patterns of a set of genes that were detected in all 3 *T. thiebautii* strains amongst the majority of the 69 samples. This consisted of a subset of 141 genes and 191 samples (out of the total 207 possible samples for 3 strains across 69 colonies). A principal coordinates (PCO) analysis indicated that many of the selected genes clustered according to their function (Fig. 6A). This grouping was confirmed using canonical analysis of principal coordinates (CAP) coupled with permutation analysis (Fig, 6B; n-permutation=999), obtaining a p=0.001 and an overall prediction accuracy of 65 %, with accuracy per group ranging between 35 % for PSII genes and 100 % for phosphate uptake, cytochrome C, chemotaxis, pili formation, and circadian rhythm, genes. This analysis revealed that the expression of genes contributing to the same overall function is not always coordinated, with some of the genes within a functional category having more similar expression patterns than others. For example, PSII genes *psb*A, *psb*E, *psb*J, and *psb*L clustered together, away from *psb*B and *psb*C which occurred together at a distance in the PCO space (Fig. 6A). The grouping of genes belonging to the same functions or operons serves as an additional confirmation that our approach correctly reflects gene expression patterns.

**Figure 6.**
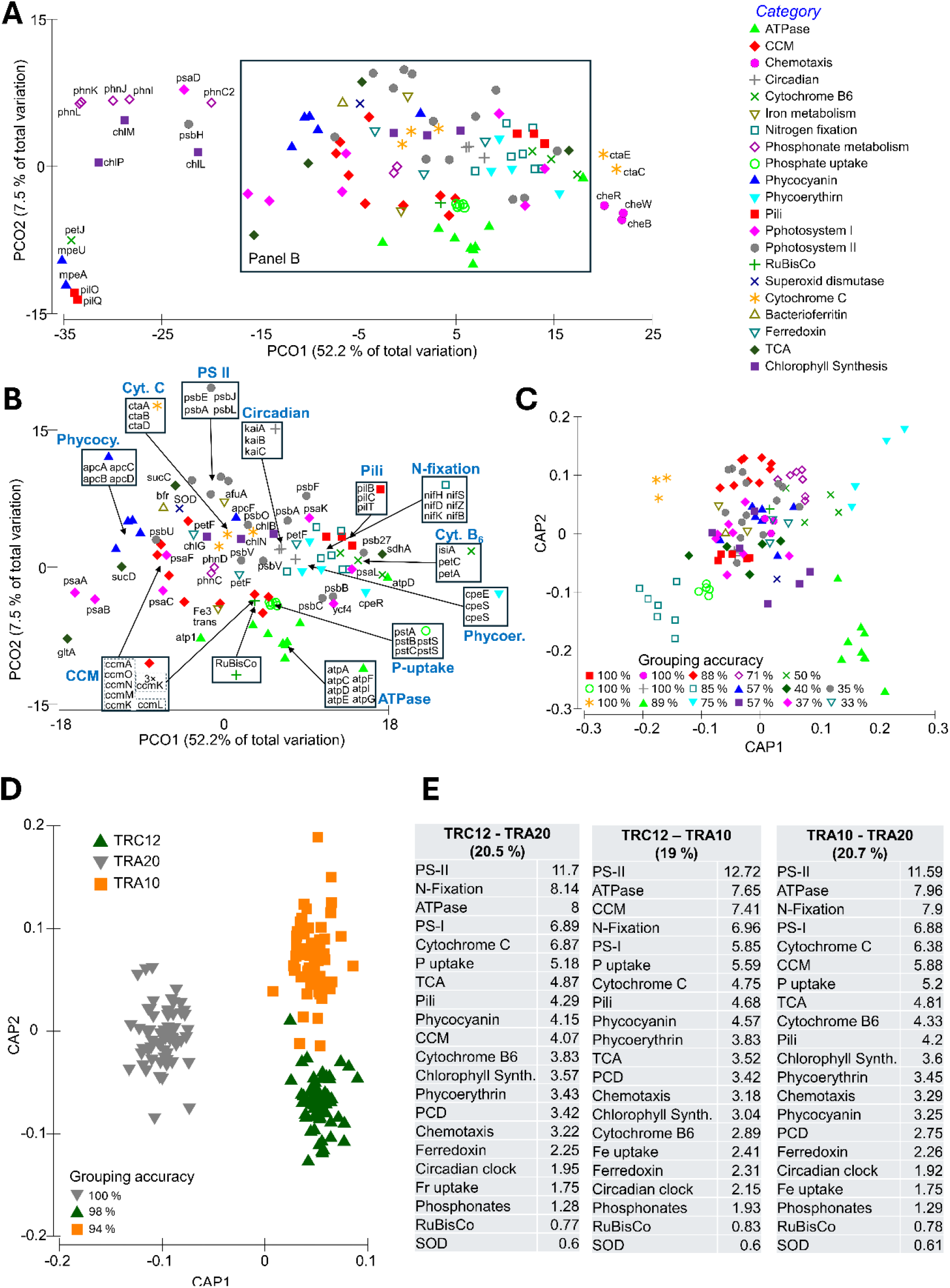
Principle COordinates analysis (PCO; A,B) and Canonical Analysis of principle Coordinates (CAP; C) of the expression patterns of the various genes revealed that certain genes belonging to similar functional categories have, as expected, similar expression profiles. The overall CAP prediction accuracy was 65% with a significant difference between functional groups (p=0.001). The prediction accuracy of the various functional groups is stated in panel C. Panel B is a focused view of the frame in panel A. CAP analysis of the different *Trichodesmium* strains (D) reveals the three strains have distinct gene expression profiles, with an overall prediction accuracy of 97 % (p=0.001). SIMPER analysis (E) reveals an average dissimilarity of *ca*. 20 % between the gene expression profiles of the strains. The contribution of different functional categories to the dissimilarity between strains is given in the figure with 100 % equaling the total dissimilarity between two strains (ca. 20%). The values per group in panel E are the sums of the individual contributions of the genes in the group.

To evaluate the difference in expression pattern between the 3 different strains (TRA10, TRA20, TRC12) we conducted a CAP analysis on the above dataset focusing on the strains rather than the genes (Fig. 6D). This revealed that the three strains have a distinct expression profile (p=0.001 and prediction accuracy of over 94 % for each of the strains). SIMPER (Similarity Percentage) analysis revealed an average dissimilarity of 20 % between the activity profiles of the strains. This dissimilarity was mainly driven by photosynthesis, ATP synthesis, N-fixation, respiration, and P uptake genes. A similar CAP analysis was conducted on all 875 common genes as identified through annotation (i.e. gene name) between the three strains, revealing an identical pattern (Fig. S3).

Using the functional analysis of the three different *Trichodesmium* strains we investigated whether the relative abundance of some of the organisms inhabiting the *Trichodesmium* colonies is correlated to the expression of specific genes by *Trichodesmium* (Table S3). The abundance of *Trichodesmium* itself, which was included in the analysis, was not correlated with any of its gene expression profiles. Out of the 233 evaluated non-cyanobacterial bacterial and archaeal genera present in at least 10 of the 69 colonies, 97 were significantly correlated with at least one gene expressed by at least one *Trichodesmium* strain. Of these genera, 26 were uniquely correlated to the activity of one strain, 17 with two strains, and 54 with all three strains.

The correlated genes were aggregated into categories to better link the different taxa to *Trichodesmium* activity. For categories with multiple genes, we required that the taxon is positively and significantly correlated (after FDR correction) to more than 3 genes from the category to state that there is a correlation to the category. According to these criteria, 10 taxa were correlated to PSII and 10 to PSI, however not all taxa were correlated with both PSI and PSII (Table S3). *Pelagibacter* sp. was associated with most photosynthesis genes. The abundance of the same strain was correlated with ATPase activity, nitrogen fixation, and phosphonate metabolism. The archaeon *Poseidonia* sp. was correlated to nitrogen fixation and so were strains of *Bacteroidia, Bdellovibrionata*, and *Alphaproteobacteria* (Table S3).

Benefiting from the transcriptomic analysis across 69 samples and the availability of 3 nearly complete *Trichodesmium* MAGs, we investigated using complete KEGG modules the correlation between the expression of different metabolic pathways in *Trichodesmium* (Fig. S4; Table S5). While many pathways were significantly and strongly correlated to others (P_adj_<=0.05), we obtained several major clusters of co-expressed pathways. One such cluster includes nitrogen-related pathways such as N fixation, nitrate assimilation, nucleotide synthesis, and the shikimate pathway. A second, larger cluster, includes basic metabolic functions in *Trichodesmium* such as PSI, Calvin cycle, NAD synthesis, ATPase, Coenzyme-A synthesis, and the citrate cycle. PSII did not cluster with any pathway and had the strongest correlation with nitrogen fixation and NAD synthesis.

## Discussion

Our current understanding of microbial metabolism in aquatic environments is mostly based on sampling and size fractionation of large volumes of water ranging between hundreds of milliliters to hundreds of liters. Nevertheless, these sampling approaches vastly exceed the size scales at which individual microorganisms experience their surrounding environment. Similarly, interactions between co-occurring organisms on organic aggregates are often concluded from the pooled average of tens to thousands of individual particles, with no confirmation that organisms co-occur on the same particle.

We present a new approach that provides coupled genomic and transcriptomic information from low-biomass samples, such as individual *Trichodesmium* colonies. This approach allows the investigation of the transcriptional activity of individual organisms whose genomes can be assembled from the obtained data, reflecting on which aspects of their metabolic potential are materialized and their specific contribution to the community’s activity. Nevertheless, in contrast to metatranscriptomics, generating a picture of the total transcriptional activity of the community is difficult.

Using this approach on nearly 70 individual *Trichodesmium* colonies we show that 1) *Trichodesmium* colonies do not harbor a core microbiome and are likely colonized by opportunistic *Bacteria* and *Archaea*; 2) Colonies harbor multiple strains of the same species, each of which has a distinct transcriptional activity profile; 3) *Trichodesmium* activity varies significantly across colonies and functions with some functions being homogenous across all colonies while other expressed by few, apparently highly active, colonies. We further confirm that *Trichodesmium* colonies harbor other filamentous and unicellular *Cyanobacteria*, as previously suggested.

OM particles in the ocean are nutrient-rich islands attracting microorganisms from the surrounding oligotrophic water ^26^ *Trichodesmium* colonies are not different. Several studies brought evidence of stochastic attachment of bacteria to OM particles, independent of particle source ^17,19,20^. In contrast, it was suggested that *Trichodesmium* colonies harbor a common core microbiome with which the nitrogen-fixing *Cyanobacteria* conducts a mutualistic relationship ^9,27^. The heterogeneous community across the 69 colonies studied here, with almost no bacterial or archaeal taxa present in all colonies, suggests there are no obligatory associations between the attached community and *Trichodesmium*. This is supported by the low number and weakness of association between colonizers and *Trichodesmium* as obtained through network analysis. The type of data used in this study originates from combined DNA and RNA. Thus, it represents a combination of sequence-frequency-based abundance (DNA) and activity (RNA). While this may affect the relative representation of certain taxa in the dataset, presence-absence data would point toward the existence of a core microbiome. However, a presence histogram (Fig. S6) clearly shows that out of 2185 identified genera, only 4 are present on all colonies and 2015 are present on less than 10 colonies.

Several different taxa are correlated with the same rudimentary *Trichodesmium* activities. While this is no clear indication of interaction or transfer of metabolites, it shows that multiple and diverse taxa are attracted to *Trichodesmium* colonies. Considering the correlations with a category of genes (e.g. PSI, PSII, N-fixation) the abundance of many taxa was positively correlated with activities such as photosynthesis or respiration, suggesting that colonizing a photosynthetically active colony was the main general attractant.

*Trichodesmium* colonies harbored multiple strains of a single *Trichodesmium* species, yet harbored other *Cyanobacteria*. During this bloom, *T. thiebautii* was the sole dominant *Trichodesmium* species detected. Previous studies in the area have shown the cooccurrence of other *Trichodesmium* species^12^. Coupled with the presence of other filamentous and unicellular *Cyanobacteria* in the colony, we propose that the mostly single-species trait results from spatial, temporal, and physiological niche separation, between *Trichodesmium* species rather than competitive interactions. However, this should be further evaluated through single-colony studies during future blooms.

Different functional categories exhibit different expression patterns across the 69 colonies. The activity of superoxide dismutase, detoxifying oxygen radicals, is homogenous across the colonies. The latter was suggested to support the activity of the oxygen-sensitive nitrogenase enzyme^28^, matching the equally homogenous activity of most genes involved in N fixation. *Trichodesmium*, like other marine cyanobacteria is known to be limited in Fe ^29^. Overall the Fe uptake category is rather heterogenous, however, this appears to be driven by two different systems. The ferrous (Fe II) iron uptake *feo*AB genes are active in a few colonies only, whereas the ferric iron (Fe III) uptake gene *fao*A and another Fe III transporter are much more homogenous. This suggests a variability in available iron sources among the colonies. Similarly, a difference in P uptake is observed. Handling of organic phosphate either via alkaline phosphatase or via the phosphonate (*phn*) genes is highly heterogeneous and active in a few colonies only. In contrast the Pi uptake *pst* genes are more homogenously active across the colonies. The heterogeneous activity of alkaline phosphatase in individual colonies could also be confirmed through direct measurements during a subsequent *Trichodesmium* bloom (Fig. S5). Similar to Fe, the P-uptake-related genes reveal that the colonies sense different P sources in their microenvironment, reacting differently one from another.

In line with the different expression patterns of different functional groups, correlation analysis of KEGG modules (Fig. S5) reveals that modules contributing to the same overall process are not necessarily co-expressed. For example, Cytochrome C oxidase and the F-type ATPase, both involved in respiration, are weakly correlated and cluster separately. Similarly, Photosystem II (PS II) is not included in any cluster and does not correlate with (PS I). This reflects the complex processes involved in the regulation and coordination of these pathways, likely involving posttranscriptional regulation and additional dependence on protein and mRNA turnover. Coupled with the advantages of single-colony studies, as revealed here, such results highlight the need to further develop low-biomass proteomics approaches as used in Held et al., 2021 ^23^ to better understand the processes that co-occur in cells.

Within each colony multiple strains of *T. thiebautii* have distinct activity profiles, suggesting complementary activity and subtle niche differentiation. Recent advances in community theory reveal that phenotypic variability among strains of the same species contributes to the success of the species in contrast to a highly specialized species lacking strain variability ^30^. Strain variability analysis revealed a much higher variability than could be resolved through genome assemblies. This suggests that the overall phenotypic (and genotypic) diversity of individual *Trichodesmium* species, and subsequently colonies, is high and may increase the ability of the entire bloom to obtain nutrients and withstand environmental stress. The lack of a fixed core microbiome suggested by our analyses does not contradict interspecies interactions with colonizing microbiota. In contrast, the specific correlation of different organisms with the abundance and activity of different *Trichodesmium* strains, suggests that the strain diversity of *Trichodesmium* expands to the associated microbiome, where these interactions are further fine-tuned through strain-specific associations with colonizing heterotrophs.

## Conclusions

Using a novel approach, we successfully resolved the transcriptional activity of different strains of *Trichodesmium thiebautii* within individual colonies. Our findings unveil a significant heterogeneity between colonies in both associated microbes and *Trichodesmium* activity. This diversity in transcriptional activity, particularly concerning nutrient uptake, underscores the highly heterogeneous nature of the colony’s microenvironment. Based on the distinct gene expression profile of each *Trichodesmium* strain we propose that *Trichodesmium* copes with the evident variable microenvironment through strain-level phenotypic diversity. Such adaptability not only refines the response of *Trichodesmium* in each colony to the environmental conditions but also appears to influence the abundance and likely activity of *Trichodesmium*-strain-specific associated heterotrophs, potentially enhancing colony fitness.

Although our sequencing depth limited the assembly of other genomes than *Trichodesmium*, our observations shed light on the dynamic interactions within *Trichodesmium* colonies and highlight the importance of microbial diversity in marine ecosystems. Our study sets the stage for future investigations exploring the effects of strain variability and strain-specific heterotrophic interaction on the *Trichodesmium* species and colony fitness.

## Acknowledgments

The fieldwork in this study was made possible through project HETRIC funded by the ASSEMBLE Plus Transnational Access program. M.B. was additionally funded through the DFG Eigene Stelle project (BI 1987/2-1). D.I. was additionally funded through the DFG Eigene Stelle project (DI 98/3-1). The authors thank Dr. Rajat Karnatak for fruitful discussions.

## Online Methods

### Sample collection

Trichodesmium colonies were collected from the Gulf of Aqaba (29.56 °N, 34.95 °E) at the Northern Red Sea during 5 consecutive days in November 2019 via morning net tows (09:00-12:00). Each tow was conducted for ∼7 min at the boat’s minimal speed (1-2 knots) by deploying a 100 μm phytoplankton net equipped with a 100 μm cod end (Aquatic Research Instrument, USA) to 10-20 m depth. The net concentrate was diluted into ∼5 L seawater to minimize stress and well-shaped puff colonies were quickly hand-picked by droppers, placed in clean Petri dishes, and washed three times with 0.22 μm filtered seawater. On each day 20 puff colonies were randomly selected and each colony was placed into a DNase and RNase-free Eppendorf (Cat. LMCT1 7B-500, Lifegene) individually with 5 μL FSW using a 20 μL pipette. Samples were immediately frozen in liquid N_2_ and kept at -80 °C until further processing.

### Nucleic Acid extraction and processing

Tubes with individual colonies were defrosted, excessive water was removed, and the colony was immersed in a lysis buffer consisting of 3 μl 1X PBS and 3 μL D2 buffer from the Repli-G single cell kit (ref. 150343, Qiagen, Germany). Cell lysis was done by four cycles of freeze/thaw, each of 1 min liquid Nitrogen and 1 min at 65 °C, followed by a 20 min incubation at 65 °C. The lysis buffer was neutralized with 3 μl of Stop solution from the Repli-G single-cell kit. Upon neutralization, 1 μL of RiboLock RNAse inhibitor (ref. EO0381, Thermo Fisher Scientific, US) was added to each tube.

The High-Capacity cDNA Reverse Transcription Kit (Cat. 4368814, Thermo Fisher Scientific) was used to generate cDNA from the RNA component of the mixed extracted nucleic acids. The final 19 μl reaction consisted of 10 μl colony extract with 1.9 μl reaction buffer, 0.75 μl dNTPs mix, 1 μl random heptamers, 1 μl reverse transcriptase, and 5.35 μl H_2_O. The reaction was incubated according to the manufacturer’s instructions for 10 min at 25 °C, 120 min at 37 °C, and 5 min at 85 °C, after which the reaction was shortly cooled down to 4 °C and immediately used for whole genome amplification (WGA). For WGA, 29 μl of Repli-G single-cell reaction buffer, and 2 μl of DNA polymerase were added to each tube, followed by an 8 h incubation at 30 °C and 3 min inactivation at 65 °C. The amplified DNA was quantified using a Quantus Fluorometer (Cat. E6150, Promega, Germany) and the QuantiFluor® ONE dsDNA System kit (Cat. E4870, Promega, Germany), after which they were frozen at -20 °C till they were shipped for sequencing.

### Sequencing

Shotgun sequencing was conducted using a NovaSeq sequencer using the S4 chip (2x150; Illumina Inc San Diego, CA, USA) after shotgun Library Prep at the Rush University Genomics and Microbiome Core Facility. Genomic DNA samples were prepared for sequencing by an initial quantification using Qubit 4 Fluorometer (Life Technologies, #Q32851, Grand Island, NY, USA). Library preparation was performed using the Illumina DNA Prep Workflow with UDI indexing (#20018705, 20027213 Illumina Inc San Diego, CA, USA) according to the manufacturer’s instructions with 50 ng template input and 5 cycles of PCR. An equal-volume pool of all libraries was then made. The pool was quantified using a Qubit DNA High Sensitivity kit (Life Technologies, #Q32851, Grand Island, NY, USA), and size distribution was assessed using an Agilent 4200 TapeStation System (Agilent Technologies, G2991AA, Santa Clara, CA, USA) using TapeStation D5000 ScreenTape, ladder and assay (Agilent Technologies, # 5067-5588, 5067-5590 and 5067-5589, Santa Clara, CA, USA). The pooled libraries were run on Illumina MiniSeq instrument using MiniSeq Reagent MO Kit, (300 cycles) (Illumina Inc San Diego, CA, USA) run for quality control and libraries balancing purposes. A new pool was made based on the MiniSeq run results, quantified same as described above and sequenced on an Illumina NovaSeq 6000 instrument (300 cycles) (Illumina Inc San Diego, CA, USA), with a 1% phiX spike-in.

The sequencing data has been deposited in the SRA under Project number: PRJNA1106210 (http://www.ncbi.nlm.nih.gov/bioproject/1106210)

### Bioinformatics

#### Sequence quality control and trimming

The quality of the raw reads was assessed using FastQC v. 0.11 (Babraham Bioinformatics). Subsequently, the raw sequence reads were quality trimmed and filtered using Trimmomatic (v. 0.39)^31^ using the command “trimmomatic PE -threads 32 <output file names> LEADING:15 TRAILING:15 SLIDINGWINDOW:4:15 MINLEN:36 HEADCROP:13”. Upon trimming, the quality of the filtered reads was assessed using FastQC.

#### Community composition analysis

The microbial community composition of the samples was assessed using CAT and RAT from the BAT/CAT/RAT package (https://github.com/MGXlab/CAT_pack)^32,33^. The latter package assigns taxonomy in a hierarchical order of reliability by mapping raw reads to assembled bins (see below), annotating contigs based on multiple identifiable protein-coding regions per contig, and lastly assigning taxonomy to unmapped reads (RAT)^33^. Microbial abundance was obtained using CoverM contig (https://github.com/wwood/CoverM), summing up the TPM values per taxon per sample. In the case of our samples, 90-99 % of the reads could be mapped back to the contigs, hence, bacterial, and archaeal taxonomy was assigned using CAT with the matching GTDB database^34^. As the GTDB database does not include eukaryotic sequences, the latter were obtained using RAT with the matching NR database.

#### MAG assembly

As the samples consist of amplified metagenomic and metatranscriptomics data multiple assembly and binning strategies were attempted. These are briefly mentioned for readers with similar data types. First, Megahit v. 1.2.9 ^35^ was used to assemble individual samples as well as a co-assembly of the entire dataset (69 samples). Subsequently, the SPAdes assembler v. 3.15.5 ^36^ was used to assemble each sample separately using either default settings, the single cell (sc) flag, or the metagenomic (meta) flag.

Binning was conducted using VAMB^37^, SemiBin2^38^, MaxBin2^39^, Metabat2 v. 2.12.12 ^40^. Contig abundance files required for the binning tools were generated by mapping the raw data to the assembled contigs with MiniMap2 ^41^, converting the SAM files to BAM using SamTools ^42^, and merging of the resulting BAM files using the jgi_summarize_bam_contig_depths script included with Meatabat2 ^40^.

The data used for further analysis was generated using the Megahit assembly of individual samples, subsequent binning with Metabat2 ^40^ and MaxBin2 ^39^, and bin refinement using MetaWrap^43^. The decision to keep these data resulted from comparing the number of bins, bin quality, and bin contamination, obtained from each assembly and binning protocol. The bins were further dereplicated using dREP v. 3.4.2 ^44^ at 99% average nucleotide identity. Trichodesmium bins were refined by removing non-cyanobacterial contigs identified using BAT ^32^.

#### MAG analysis

The quality of the obtained bins was analyzed using Checkm^45^ and Checkm2^46^. Taxonomy was assigned to bins using GTDB-tk v. 2.3.2 ^25^ with database r214. Before quantifying the abundance of different *Trichodesmium* strains (dereplicated at 99 % average nucleotide identity) were quantified using CoverM v. 0.6.1 (https://github.com/wwood/CoverM) and strain diversity analysis with InStrain^24^ v. 1.3.1.

Annotation of the *Trichodesmium* MAGs was done initially using Prokka v. 1.14.5^47^. Subsequently, all protein-coding open reading frames called using the Prokka pipeline were further annotated using the EggNog mapper (emapper v. 2.1.12)^48,49^, as well as against the KEGG^50^ database using the BlastKoala^51^ and KofamKoala ^52^.

#### Gene expression quantification

The sequence data contains sequences originating from both genomic DNA and RNA. The subsequent calculations consider a fixed ratio between gene copy numbers in a given genome and therefore, any deviation from that ratio would come from RNA data.

First, transcripts per kilobase million (TPM) values were calculated for the genes of each strain for each sample using Salmon v. 1.10.1 ^53^. Subsequently, a relative gene expression value was generated by dividing the TPM values of each protein by that of the *rpo*B gene. The latter is a housekeeping gene for which minimal variation in expression is expected. The basal ratio between the expression of each gene and that of *rpo*B is unknown therefore, for comparison purposes, the values for each gene were scaled separately for each *Trichodesmium* strain between 0 and 1, where 1 was the sample (colony) with the highest value for that particular gene. Zero values were treated as missing data rather than an absolute lack of expression.

We used the *Trichodesmium* transcriptome study published by Cerdan-Garcia ^15^ to check whether normalizing data to *rpo*B values biases the results. The exact data used in the paper was courteously provided by the authors and normalized to their rpoB values. Subsequently, a differential expression analysis was done using the online tool iDEP v.1.13 (http://bioinformatics.sdstate.edu/idep11/;^54^. A Principal Component Analysis (PCOA) of the samples was conducted using Primer6 (v.6.1.1) + PERMANOVA package (v.1.0.1, Primer-E, Quest Research Limited, Auckland, New Zealand).

#### Detection of alkaline phosphatase activity (APA)

Alkaline phosphatase activity (APA) of Red Sea *Trichodesmium* colonies was quantified by tracking the enzymatic cleavage of 4-methylumbelliferyl phosphate (4-MUP) (Sigma; Cat#M8883-250MG). The assays were performed using either bulk colonies (3-5 colonies per sample) or individual colonies and daily-prepared microwaved seawater (MFSW) with the presence of 10 mM Tris buffer (pH=8.2) and 5 μg mL_-1_ 4-MUP at 25°C in the dark. Fluorescence of samples were measured at two intervals, t=0 and t=1 hour, with the colony were taken out from the cuvette to avoid fluctuations. The fluorescence was measured using a fluorescence spectrophotometer (Cary Eclipse, Agilent, USA) at an excitation wavelength of 385 nm and emission wavelength of 440 nm ^55^. APA was calculated from the linear regression of fluorescence against time, denoted in the unit of fluorescence per minute and it was then converted to nmol 4-MUP colony^-1^ h^-1^ using a calibration curve. The blank was measured daily in the same manner using MFSW and was then subtracted from samples. Individual colonies were imaged before APA measurements using a stereoscope (Nikon, SMZ745) equipped with the Dino-Eye eyepiece camera and colony size was determined using DinoCapture 2.0 software.

#### Further analyses

Network analysis was conducted using the community datasets obtained with CAT ^32^ using the NetCommi R package ^56^ with SparCC ^57^as the network construction approach. The resulting networks were visualized using Gephi.

To test for correlation between expression patterns of different KEGG modules, a module completeness analysis was done with all genes that received a KO during the annotation step. For modules for which one block of missing and were known to occur in other Trichodesmium or Cyanobacteria, the missing gene was searched by name in the EggNog annotation and if found, assigned a KO. The correlation analysis was conducted using the corr.test function of the Psych R package (https://personality-project.org/r/psych/) using Pearson correlation and Benjamini Hochberg adjustment of the P values. The correlation plot was done using the corrplot R package (https://cran.r-project.org/web/packages/corrplot/vignettes/corrplot-intro.html), with KEGG module order being derived from a bootstrapped clustering (n=10,000) with the pvclust R package (https://github.com/shimo-lab/pvclust) using the Ward.D2 clustering method ^58^. The Jensen-Shanno metric was calculated using the distance.jensenshannon function from the scipy.spatial Python library, comparing the expression pattern of each gene to a homogenous vector of 69 data points each equalling 1. Missing data was omitted and the resulting vector was compared to a homogenous vector of the same size. Shannon Evenness was calculated by dividing the Shannon entropy per gene by the maximum entropy in the system.

Principal coordinate analysis of the gene expression data (ratios to *rpo*B) per sample or per gene (see results) and subsequent canonical analysis of principal coordinates were done on selected genes representing major pathways using Primer6 (v.6.1.1) + PERMANOVA package (v.1.0.1, Primer-E, Quest Research Limited, Auckland, New Zealand).

## Notes

### Competing Interest Statement

The authors have declared no competing interest.

