## Supplementary tables and figures for "Phenotypic variability between closely related *Trichodesmium* strains as a means to making the most out of heterogeneous microenvironments"

**Table S1.** Average Nucleotide Identity (ANI) between Trichodesmium and Okeania Genomes (MAGs). Trichodesmium strains in marked red were obtained in this study and strains marked in blue were obtained in other studies from the same location.

| ANI | Okeania_GCA_003838195.1 | Okeania_GCA_003838225.1 | Okeania_GCA_010672015.1 | Okeania_GCA_010672085.1 | Okeania_GCA_010672215.1 | Okeania_GCA_010672295.1 | Okeania_GCA_010672305.1 | Okeania_GCA_010672385.1 | Okeania_GCA_010672395.1 | Okeania_GCA_010672455.1 | Okeania_GCA_010672505.1 | Okeania_GCA_010672525.1 | Okeania_GCA_010672585.1 | Okeania_GCA_010672625.1 | Okeania_GCA_010672645.1 | Okeania_GCA_010672685.1 | Okeania_GCA_010672695.1 | Okeania_GCA_010672745.1 | Okeania_GCA_010692435.1 | Okeania_GCA_010692535.1 | Okeania_GCA_010692555.1 | Okeania_GCA_010692585.1 | Okeania_GCA_010692625.1 | Okeania_GCA_010692685.1 | Okeania_GCA_014132355.1 | Okeania_GCA_035285525.1 |
|---|---|---|---|---|---|---|---|---|---|---|---|---|---|---|---|---|---|---|---|---|---|---|---|---|---|---|
| **TRA10_S76.bins.1** | 80.9 | 81.0 | 80.5 | 83.9 | 79.8 | 81.0 | 81.0 | 83.8 | 79.6 | 80.7 | 80.7 | 80.7 | 80.6 | 81.0 | 81.0 | 80.9 | 80.9 | 81.0 | 80.4 | 80.3 | 80.8 | 80.5 | 80.8 | 81.0 | 80.7 | 81.6 |
| **TRA20_S83.bins.1** | 81.0 | 81.0 | 80.4 | 83.8 | 79.8 | 81.0 | 81.0 | 83.8 | 79.4 | 80.8 | 80.7 | 80.7 | 80.7 | 80.9 | 81.0 | 80.9 | 80.9 | 81.0 | 80.3 | 80.4 | 80.8 | 80.5 | 80.8 | 81.0 | 80.8 | 81.4 |
| **TRC12_S101.bins.1** | 81.0 | 81.0 | 80.6 | 83.9 | 79.8 | 80.9 | 80.9 | 83.9 | 79.9 | 80.8 | 80.7 | 80.7 | 80.7 | 81.0 | 81.1 | 81.0 | 81.0 | 80.9 | 80.5 | 80.5 | 80.8 | 80.6 | 80.8 | 80.9 | 80.8 | 81.6 |
| Tricho_GCA_000963755.2 | 80.7 | 80.8 | 80.5 | 83.8 | 80.0 | 80.7 | 80.7 | 83.8 | 79.5 | 80.6 | 80.5 | 80.7 | 80.7 | 80.8 | 80.7 | 80.8 | 80.8 | 80.8 | 80.4 | 80.5 | 80.7 | 80.3 | 80.8 | 80.8 | 80.6 | 81.8 |
| Tricho_GCA_000987385.1 | 80.8 | 80.7 | 80.0 | 84.1 | 79.6 | 80.7 | 80.7 | 83.7 | 79.2 | 80.7 | 80.4 | 80.7 | 80.5 | 80.7 | 80.7 | 80.7 | 80.8 | 80.7 | 80.3 | 80.4 | 80.6 | 80.1 | 80.9 | 80.7 | 80.6 | 81.5 |
| Tricho_GCA_018457135.1 | 80.8 | 80.9 | 80.6 | 83.7 | 80.0 | 80.8 | 80.8 | 83.7 | 79.6 | 80.6 | 80.6 | 80.6 | 80.6 | 80.8 | 80.9 | 80.8 | 80.8 | 80.8 | 80.4 | 80.5 | 80.6 | 80.4 | 80.9 | 80.9 | 80.6 | 81.8 |
| Tricho_GCA_022448615.1 | 80.9 | 80.9 | 80.5 | 83.8 | 80.0 | 80.9 | 80.9 | 83.8 | 79.4 | 80.6 | 80.7 | 80.6 | 80.7 | 80.9 | 80.9 | 80.8 | 80.8 | 80.8 | 80.5 | 80.5 | 80.6 | 80.4 | 80.9 | 80.9 | 80.6 | 81.8 |
| Tricho_GCA_022822685.1 | 80.1 | 80.1 | 79.4 | 79.6 | 79.4 | 80.2 | 80.2 | 80.1 | 78.5 | 80.0 | 79.8 | 80.2 | 80.1 | 80.1 | 79.9 | 80.2 | 80.2 | 80.1 | 79.9 | 79.4 | 79.9 | 79.7 | 80.1 | 80.2 | 79.8 | 78.7 |
| **Tricho_GCA_023356515.1** | 80.7 | 80.7 | 79.7 | 84.1 | 79.5 | 80.7 | 80.7 | 84.0 | 79.0 | 80.6 | 80.3 | 80.6 | 80.7 | 80.7 | 80.5 | 80.7 | 80.7 | 80.7 | 80.2 | 80.0 | 80.5 | 79.9 | 80.7 | 80.7 | 80.4 | 81.7 |
| **Tricho_GCA_023356535.1** | 80.9 | 80.9 | 80.3 | 84.5 | 79.9 | 80.9 | 80.9 | 84.4 | 79.5 | 80.9 | 80.6 | 80.9 | 80.9 | 80.9 | 80.9 | 80.9 | 80.9 | 80.9 | 80.7 | 80.4 | 80.8 | 80.1 | 80.9 | 80.9 | 80.7 | 81.9 |
| **Tricho_GCA_023356555.1** | 80.8 | 80.9 | 80.6 | 83.8 | 80.1 | 80.9 | 80.8 | 83.9 | 79.7 | 80.7 | 80.6 | 80.7 | 80.7 | 80.8 | 80.8 | 80.8 | 80.8 | 80.9 | 80.5 | 80.5 | 80.7 | 80.4 | 80.8 | 80.8 | 80.5 | 81.8 |
| **Tricho_GCA_023356605.1** | 80.9 | 81.0 | 80.5 | 84.1 | 79.7 | 80.9 | 81.0 | 83.9 | 79.6 | 80.8 | 80.7 | 80.7 | 80.7 | 80.9 | 80.9 | 80.9 | 80.9 | 80.9 | 80.5 | 80.4 | 80.8 | 80.5 | 80.9 | 81.0 | 80.9 | 81.6 |
| Tricho_GCA_913057915.1 | 80.7 | 80.8 | 80.6 | 83.8 | 80.1 | 80.8 | 80.9 | 83.8 | 79.5 | 80.6 | 80.6 | 80.6 | 80.6 | 80.8 | 80.8 | 80.8 | 80.8 | 80.8 | 80.4 | 80.5 | 80.7 | 80.4 | 80.8 | 80.9 | 80.6 | 81.8 |

**Table S2.** Taxa significantly correlated with *Trichodesmium* strains abundance.

| **Taxonomy** | **Tricho** | **R** | **p** | **P adj** | **# Colonies present** |
|---|---|---|---|---|---|
| d_Bacteria; p_Bacteroidota; c_Bacteroidia; o_Flavobacteriales; f_Flavobacteriaceae; g_Gilvibacter | TRA10 | 0.58 | 1.59E-07 | 4.94E-05 | 69 |
| d_Bacteria; p_Pseudomonadota; c_Alphaproteobacteria; o_Pelagibacterales; f_Pelagibacteraceae; g_Fonsibacter | TRA10 | 0.43 | 0.000178 | 0.020683 | 21 |
| d_Bacteria; p_Bacteroidota; c_Bacteroidia; o_NS11-12g; f_UBA9320; g_UBA9320 | TRC12 | 0.59 | 5.63E-08 | 2.62E-05 | 33 |
| d_Bacteria; p_Bacteroidota; c_Bacteroidia; o_Flavobacteriales; f_Flavobacteriaceae; g_Polaribacter | TRC12 | 0.45 | 9.38E-05 | 0.014056 | 33 |
| d_Bacteria; p_Bacteroidota; c_Bacteroidia; o_Flavobacteriales; f_Crocinitomicaceae; g_UBA4466 | TRC12 | 0.63 | 4.63E-09 | 4.31E-06 | 24 |
| d_Bacteria; p_Pseudomonadota; c_Alphaproteobacteria; o_TMED127; f_TMED127; g_GCA-2691145 | TRC12 | 0.44 | 0.000106 | 0.014056 | 17 |
| d_Bacteria; p_Marinisomatota; c_Marinisomatia; o_SCGC-AAA003-L08; f_GCA-002704045; g_GCA-002704045 | TRC12 | 0.47 | 3.17E-05 | 0.005912 | 15 |
| d_Bacteria; p_Pseudomonadota; c_Gammaproteobacteria; o_SAR86; f_D2472; g_MED-G82 | TRC12 | 0.54 | 1.05E-06 | 0.000245 | 10 |

**Table S3.** Significantly correlated taxa with the transcriptional activity of the three *Trichodesmium* strains organized according to functional categories. The numbers in each cell represent the number of correlated genes from that category.

| | | Bacteria; Pseudomonadota; Fonsibacter | Bacteria; Bacteroidota; 2-12-FULL-35-15 | Bacteria; Verrucomicrobiota; GCA-2699365 | Bacteria; Marinisomatota; GCA-002704045 | Archaea; Thermoproteota; Nitrosopelagicus | Bacteria; Bacteroidota; Kordia | Bacteria; Pseudomonadota; Cellvibrio | Bacteria; Bdellovibrionota; UBA6935 | Bacteria; Marinisomatota; CACEUX01 | Bacteria; Pseudomonadota; MED-G82 | Bacteria; Pseudomonadota; GCA-2691145 | Bacteria; Bacteroidota; UBA9320 | Bacteria; Bacteroidota; Polaribacter | Bacteria; Bacteroidota; UBA4466 | Bacteria; Pseudomonadota; Hyphobacterium | Bacteria; Bacteroidota; UWMA-0277 | Bacteria; SAR324; Arctic96AD-7 | Bacteria; Pseudomonadota; GCA-2732015 | Archaea; Thermoplasmatota; Poseidonia | Bacteria; Bacteroidota; Winogradskyella | Bacteria; Bacteroidota; GCA-2711565 | Bacteria; Actinomycetota; Streptomyces | Bacteria; Pseudomonadota; VGCX01 | Archaea; Nanoarchaeota; ARS1285 | Bacteria; Bacteroidota; Altibacter | Bacteria; Pseudomonadota; GCA-2724185 |
|---|---|---|---|---|---|---|---|---|---|---|---|---|---|---|---|---|---|---|---|---|---|---|---|---|---|---|---|
| PSI | TRC12 | 5 | 6 | 5 | 3 | 5 | 4 | 4 | 3 | 4 | 2 | 2 | 2 | 2 | 2 | 2 | 1 | 2 | 1 | 1 | 0 | 0 | 0 | 0 | 0 | 0 | 0 |
| | TRA20 | 6 | 0 | 0 | 4 | 0 | 0 | 0 | 2 | 0 | 3 | 3 | 3 | 3 | 2 | 2 | 3 | 0 | 2 | 1 | 0 | 0 | 0 | 0 | 0 | 0 | 0 |
| | TRA10 | 4 | 6 | 5 | 3 | 4 | 4 | 4 | 3 | 4 | 2 | 2 | 2 | 2 | 2 | 2 | 1 | 2 | 1 | 1 | 0 | 0 | 0 | 0 | 0 | 0 | 0 |
| PSII | TRC12 | 9 | 6 | 6 | 2 | 4 | 4 | 4 | 2 | 2 | 1 | 0 | 0 | 0 | 1 | 1 | 0 | 1 | 0 | 2 | 4 | 4 | 4 | 3 | 2 | 1 | 1 |
| | TRA20 | 10 | 0 | 0 | 1 | 0 | 0 | 0 | 1 | 0 | 0 | 0 | 0 | 0 | 0 | 0 | 1 | 0 | 0 | 4 | 4 | 4 | 3 | 3 | 0 | 0 | 0 |
| | TRA10 | 8 | 6 | 6 | 2 | 5 | 5 | 4 | 1 | 2 | 1 | 0 | 0 | 0 | 1 | 1 | 0 | 1 | 0 | 2 | 5 | 4 | 3 | 3 | 1 | 1 | 1 |
| RUBISC | TRC12 | 0 | 0 | 0 | 1 | 0 | 0 | 0 | 0 | 0 | 1 | 1 | 1 | 0 | 0 | 0 | 0 | 0 | 0 | 0 | 0 | 0 | 0 | 0 | 0 | 0 | 0 |
| | TRA20 | 0 | 0 | 0 | 1 | 0 | 0 | 0 | 0 | 0 | 1 | 1 | 0 | 0 | 0 | 0 | 0 | 0 | 0 | 0 | 0 | 0 | 0 | 0 | 0 | 0 | 0 |
| | TRA10 | 0 | 0 | 0 | 1 | 0 | 0 | 0 | 0 | 0 | 1 | 1 | 1 | 0 | 0 | 0 | 0 | 0 | 0 | 0 | 0 | 0 | 0 | 0 | 0 | 0 | 0 |
| ATP | TRC12 | 0 | 2 | 1 | 7 | 1 | 1 | 1 | 2 | 1 | 7 | 7 | 6 | 6 | 2 | 0 | 3 | 0 | 1 | 0 | 0 | 0 | 0 | 0 | 0 | 0 | 0 |
| | TRA20 | 0 | 0 | 0 | 6 | 0 | 0 | 0 | 1 | 0 | 6 | 6 | 4 | 4 | 0 | 0 | 3 | 0 | 1 | 0 | 0 | 0 | 0 | 0 | 0 | 0 | 0 |
| | TRA10 | 0 | 2 | 1 | 7 | 1 | 1 | 1 | 1 | 1 | 7 | 7 | 6 | 6 | 1 | 0 | 4 | 0 | 1 | 0 | 0 | 0 | 0 | 0 | 0 | 0 | 0 |
| ATPase | TRC12 | 4 | 4 | 3 | 0 | 3 | 3 | 3 | 0 | 3 | 0 | 0 | 0 | 0 | 0 | 0 | 0 | 3 | 0 | 0 | 3 | 3 | 3 | 1 | 0 | 0 | 0 |
| | TRA20 | 3 | 0 | 0 | 0 | 0 | 0 | 0 | 0 | 0 | 0 | 0 | 0 | 0 | 0 | 0 | 0 | 0 | 0 | 0 | 4 | 4 | 3 | 2 | 0 | 0 | 0 |
| | TRA10 | 3 | 2 | 1 | 0 | 1 | 1 | 1 | 0 | 1 | 0 | 0 | 0 | 0 | 0 | 0 | 0 | 1 | 0 | 0 | 4 | 4 | 3 | 1 | 0 | 0 | 0 |
| CCM | TRC12 | 2 | 1 | 1 | 5 | 1 | 0 | 1 | 3 | 0 | 5 | 5 | 2 | 0 | 1 | 4 | 0 | 0 | 0 | 0 | 6 | 5 | 5 | 4 | 6 | 1 | 0 |
| | TRA20 | 4 | 0 | 0 | 3 | 0 | 0 | 0 | 3 | 0 | 4 | 4 | 1 | 0 | 1 | 3 | 0 | 0 | 0 | 0 | 7 | 7 | 5 | 5 | 5 | 0 | 0 |
| | TRA10 | 2 | 1 | 1 | 5 | 1 | 0 | 0 | 3 | 0 | 5 | 5 | 2 | 0 | 1 | 3 | 0 | 0 | 0 | 0 | 6 | 6 | 5 | 4 | 6 | 1 | 0 |
| NFIX | TRC12 | 7 | 3 | 3 | 3 | 2 | 3 | 2 | 1 | 2 | 3 | 2 | 2 | 2 | 0 | 0 | 1 | 2 | 0 | 6 | 2 | 2 | 2 | 0 | 0 | 0 | 0 |
| | TRA20 | 6 | 0 | 0 | 1 | 0 | 0 | 0 | 1 | 0 | 1 | 1 | 0 | 0 | 0 | 0 | 0 | 0 | 0 | 5 | 1 | 1 | 1 | 1 | 0 | 0 | 0 |
| | TRA10 | 7 | 3 | 2 | 4 | 2 | 2 | 2 | 2 | 2 | 4 | 4 | 3 | 3 | 0 | 0 | 2 | 2 | 0 | 6 | 2 | 2 | 1 | 1 | 0 | 0 | 0 |
| CYT | TRC12 | 2 | 1 | 1 | 6 | 0 | 1 | 0 | 6 | 0 | 6 | 6 | 6 | 5 | 6 | 2 | 5 | 0 | 1 | 0 | 2 | 2 | 1 | 1 | 1 | 1 | 1 |
| | TRA20 | 1 | 0 | 0 | 6 | 0 | 0 | 0 | 5 | 0 | 6 | 6 | 6 | 6 | 6 | 2 | 6 | 0 | 1 | 0 | 3 | 3 | 2 | 2 | 1 | 1 | 1 |
| | TRA10 | 2 | 1 | 0 | 6 | 0 | 1 | 0 | 6 | 0 | 6 | 6 | 6 | 5 | 6 | 2 | 5 | 0 | 1 | 0 | 2 | 2 | 1 | 1 | 1 | 1 | 1 |
| PHOS | TRC12 | 2 | 1 | 1 | 6 | 0 | 1 | 0 | 6 | 0 | 6 | 6 | 6 | 5 | 6 | 2 | 5 | 0 | 1 | 0 | 2 | 2 | 1 | 1 | 1 | 1 | 1 |
| | TRA20 | 1 | 0 | 0 | 6 | 0 | 0 | 0 | 5 | 0 | 6 | 6 | 6 | 6 | 6 | 2 | 6 | 0 | 1 | 0 | 3 | 3 | 2 | 2 | 1 | 1 | 1 |
| | TRA10 | 2 | 1 | 0 | 6 | 0 | 1 | 0 | 6 | 0 | 6 | 6 | 6 | 5 | 6 | 2 | 5 | 0 | 1 | 0 | 2 | 2 | 1 | 1 | 1 | 1 | 1 |
| PHN | TRC12 | 5 | 1 | 0 | 1 | 0 | 0 | 0 | 5 | 0 | 1 | 0 | 0 | 0 | 0 | 5 | 0 | 0 | 0 | 0 | 0 | 0 | 0 | 0 | 5 | 5 | 5 |
| | TRA20 | 5 | 0 | 0 | 0 | 0 | 0 | 0 | 5 | 0 | 0 | 0 | 0 | 0 | 0 | 5 | 0 | 0 | 0 | 0 | 0 | 0 | 0 | 0 | 5 | 5 | 5 |
| | TRA10 | 5 | 1 | 0 | 1 | 0 | 0 | 0 | 5 | 0 | 1 | 0 | 0 | 0 | 0 | 5 | 0 | 0 | 0 | 0 | 0 | 0 | 0 | 0 | 5 | 5 | 5 |
| IRON | TRC12 | 1 | 0 | 0 | 3 | 0 | 0 | 0 | 3 | 0 | 3 | 3 | 3 | 3 | 3 | 0 | 3 | 0 | 2 | 0 | 1 | 1 | 1 | 1 | 0 | 0 | 0 |
| | TRA20 | 1 | 0 | 0 | 2 | 0 | 0 | 0 | 1 | 0 | 2 | 2 | 2 | 2 | 2 | 0 | 2 | 0 | 1 | 0 | 0 | 0 | 0 | 0 | 0 | 0 | 0 |
| | TRA10 | 1 | 0 | 0 | 3 | 0 | 0 | 0 | 3 | 0 | 3 | 3 | 3 | 3 | 3 | 0 | 3 | 0 | 2 | 0 | 0 | 0 | 0 | 0 | 0 | 0 | 0 |
| PhycoC | TRC12 | 2 | 2 | 1 | 2 | 1 | 1 | 1 | 3 | 1 | 2 | 1 | 1 | 1 | 1 | 3 | 1 | 0 | 0 | 1 | 5 | 5 | 5 | 2 | 3 | 3 | 3 |
| | TRA20 | 2 | 0 | 0 | 2 | 0 | 0 | 0 | 2 | 0 | 2 | 2 | 2 | 2 | 2 | 2 | 2 | 0 | 2 | 0 | 4 | 4 | 4 | 3 | 2 | 2 | 1 |
| | TRA10 | 2 | 2 | 1 | 2 | 1 | 1 | 1 | 3 | 1 | 2 | 1 | 1 | 1 | 1 | 3 | 1 | 0 | 0 | 1 | 5 | 5 | 5 | 2 | 3 | 3 | 2 |
| PhycoE | TRC12 | 0 | 2 | 3 | 8 | 2 | 2 | 2 | 7 | 1 | 8 | 8 | 8 | 7 | 8 | 0 | 7 | 0 | 7 | 0 | 0 | 0 | 0 | 0 | 0 | 0 | 0 |
| | TRA20 | 0 | 0 | 0 | 1 | 0 | 0 | 0 | 1 | 0 | 1 | 1 | 1 | 1 | 1 | 0 | 1 | 0 | 1 | 0 | 0 | 0 | 0 | 0 | 0 | 0 | 0 |
| | TRA10 | 0 | 3 | 3 | 9 | 2 | 2 | 1 | 8 | 1 | 9 | 9 | 9 | 8 | 9 | 0 | 8 | 0 | 7 | 0 | 0 | 0 | 0 | 0 | 0 | 0 | 0 |
| CASP | TRC12 | 2 | 0 | 0 | 1 | 0 | 0 | 0 | 1 | 0 | 1 | 1 | 1 | 1 | 1 | 1 | 1 | 0 | 1 | 0 | 2 | 2 | 1 | 0 | 0 | 0 | 0 |
| | TRA20 | 2 | 0 | 0 | 0 | 0 | 0 | 0 | 0 | 0 | 0 | 0 | 0 | 0 | 0 | 0 | 0 | 0 | 0 | 0 | 2 | 2 | 2 | 1 | 0 | 0 | 0 |
| | TRA10 | 3 | 0 | 0 | 1 | 0 | 0 | 0 | 1 | 0 | 1 | 1 | 1 | 1 | 1 | 1 | 1 | 0 | 1 | 1 | 2 | 2 | 1 | 0 | 0 | 0 | 0 |
| SOD | TRC12 | 0 | 0 | 0 | 0 | 0 | 0 | 0 | 0 | 0 | 0 | 0 | 0 | 0 | 0 | 0 | 0 | 0 | 0 | 0 | 0 | 0 | 0 | 0 | 0 | 0 | 0 |
| | TRA20 | 0 | 0 | 0 | 0 | 0 | 0 | 0 | 0 | 0 | 0 | 0 | 0 | 0 | 0 | 0 | 0 | 0 | 0 | 0 | 0 | 0 | 0 | 0 | 0 | 0 | 0 |
| | TRA10 | 0 | 0 | 0 | 0 | 0 | 0 | 0 | 0 | 0 | 0 | 0 | 0 | 0 | 0 | 0 | 0 | 0 | 0 | 0 | 1 | 1 | 1 | 1 | 0 | 0 | 0 |
| PIL | TRC12 | 6 | 1 | 2 | 4 | 1 | 1 | 1 | 4 | 1 | 4 | 4 | 4 | 4 | 4 | 0 | 4 | 0 | 4 | 2 | 1 | 1 | 1 | 1 | 0 | 0 | 0 |
| | TRA20 | 6 | 0 | 0 | 2 | 0 | 0 | 0 | 3 | 0 | 2 | 2 | 2 | 2 | 3 | 1 | 2 | 0 | 2 | 4 | 1 | 1 | 1 | 1 | 1 | 1 | 1 |
| | TRA10 | 6 | 1 | 2 | 3 | 1 | 1 | 1 | 3 | 1 | 3 | 3 | 3 | 3 | 3 | 0 | 3 | 0 | 3 | 2 | 1 | 1 | 1 | 1 | 0 | 0 | 0 |
| CHEMC | TRC12 | 1 | 3 | 1 | 0 | 1 | 1 | 1 | 0 | 1 | 0 | 0 | 0 | 0 | 0 | 1 | 0 | 1 | 0 | 0 | 1 | 1 | 1 | 1 | 1 | 1 | 0 |
| | TRA20 | 1 | 0 | 0 | 0 | 0 | 0 | 0 | 0 | 0 | 0 | 0 | 0 | 0 | 0 | 1 | 0 | 0 | 0 | 0 | 1 | 1 | 1 | 1 | 1 | 1 | 1 |
| | TRA10 | 0 | 1 | 0 | 0 | 0 | 0 | 0 | 0 | 0 | 0 | 0 | 0 | 0 | 0 | 0 | 0 | 0 | 0 | 0 | 1 | 1 | 1 | 1 | 0 | 0 | 0 |
| DPS | TRC12 | 1 | 0 | 0 | 0 | 0 | 0 | 0 | 0 | 0 | 0 | 0 | 0 | 0 | 0 | 0 | 0 | 0 | 0 | 0 | 0 | 0 | 0 | 0 | 0 | 0 | 0 |
| | TRA20 | 1 | 0 | 0 | 0 | 0 | 0 | 0 | 0 | 0 | 0 | 0 | 0 | 0 | 0 | 0 | 0 | 0 | 0 | 0 | 0 | 0 | 0 | 0 | 0 | 0 | 0 |
| | TRA10 | 1 | 0 | 0 | 0 | 0 | 0 | 0 | 0 | 0 | 0 | 0 | 0 | 0 | 0 | 0 | 0 | 0 | 0 | 0 | 0 | 0 | 0 | 0 | 0 | 0 | 0 |
| GLTA | TRC12 | 0 | 0 | 0 | 1 | 0 | 0 | 0 | 1 | 0 | 1 | 1 | 1 | 1 | 1 | 0 | 1 | 0 | 1 | 0 | 0 | 0 | 0 | 0 | 0 | 0 | 0 |
| | TRA20 | 0 | 0 | 0 | 1 | 0 | 0 | 0 | 1 | 0 | 1 | 1 | 1 | 1 | 1 | 0 | 1 | 0 | 0 | 0 | 0 | 0 | 0 | 0 | 0 | 0 | 0 |
| | TRA10 | 0 | 0 | 0 | 1 | 0 | 0 | 0 | 1 | 0 | 1 | 1 | 1 | 1 | 1 | 0 | 1 | 0 | 1 | 0 | 0 | 0 | 0 | 0 | 0 | 0 | 0 |

**Table S4**. Correlation between transcriptional activity summed up into KEGG modules. This data is the basis for Figure 6.

| Module1 | Module2 | R | P | Padj |
|---|---|---|---|---|
| M00001 | M00004 | 0.98 | 2.74E-51 | 1.21E-49 |
| M00001 | M00010 | 0.77 | 1.35E-14 | 6.31E-14 |
| M00001 | M00015 | 0.62 | 1.50E-08 | 3.84E-08 |
| M00001 | M00018 | 0.59 | 1.02E-07 | 2.38E-07 |
| M00001 | M00019 | 0.38 | 0.001258 | 0.001834 |
| M00001 | M00021 | 0.37 | 0.001749 | 0.00249 |
| M00001 | M00023 | 0.43 | 0.000206 | 0.000338 |
| M00001 | M00024 | 0.79 | 1.00E-15 | 5.28E-15 |
| M00001 | M00028 | 0.35 | 0.003509 | 0.004778 |
| M00001 | M00048 | 0.35 | 0.003005 | 0.004133 |
| M00001 | M00049 | 0.37 | 0.001576 | 0.002261 |
| M00001 | M00050 | 0.36 | 0.002277 | 0.003177 |
| M00001 | M00052 | 0.29 | 0.014724 | 0.018475 |
| M00001 | M00083 | 0.65 | 1.40E-09 | 3.90E-09 |
| M00001 | M00086 | 0.30 | 0.011486 | 0.014716 |
| M00001 | M00096 | 0.88 | 1.53E-23 | 1.51E-22 |
| M00001 | M00097 | 0.46 | 7.97E-05 | 0.000139 |
| M00001 | M00112 | 0.29 | 0.016962 | 0.021011 |
| M00001 | M00115 | 0.78 | 2.74E-15 | 1.38E-14 |
| M00001 | M00119 | 0.38 | 0.001296 | 0.001886 |
| M00001 | M00120 | 0.91 | 6.67E-28 | 1.00E-26 |
| M00001 | M00121 | 0.56 | 4.58E-07 | 1.02E-06 |
| M00001 | M00133 | 0.92 | 1.10E-28 | 1.76E-27 |
| M00001 | M00135 | 0.36 | 0.002054 | 0.00288 |
| M00001 | M00145 | 0.80 | 1.28E-16 | 7.43E-16 |
| M00001 | M00155 | 0.38 | 0.001211 | 0.001771 |
| M00001 | M00157 | 0.56 | 4.25E-07 | 9.57E-07 |
| M00001 | M00163 | 0.93 | 3.10E-30 | 5.26E-29 |
| M00001 | M00165 | 0.95 | 8.76E-36 | 2.16E-34 |
| M00001 | M00175 | 0.46 | 6.43E-05 | 0.000114 |
| M00001 | M00176 | 0.48 | 3.25E-05 | 5.95E-05 |
| M00001 | M00307 | 0.40 | 0.000658 | 0.000996 |
| M00001 | M00432 | 0.88 | 7.67E-23 | 7.12E-22 |
| M00001 | M00527 | 0.37 | 0.001538 | 0.002218 |
| M00001 | M00549 | 0.29 | 0.016827 | 0.020862 |
| M00001 | M00621 | 0.30 | 0.01312 | 0.016591 |
| M00001 | M00793 | 0.32 | 0.006687 | 0.008792 |
| M00001 | M00842 | 0.37 | 0.001893 | 0.002677 |
| M00001 | M00843 | 0.36 | 0.002128 | 0.002978 |
| M00001 | M00844 | 0.73 | 9.13E-13 | 3.49E-12 |
| M00001 | M00854 | 0.97 | 5.77E-45 | 2.16E-43 |
| M00001 | M00880 | 0.35 | 0.003229 | 0.004429 |
| M00001 | M00881 | 0.40 | 0.000577 | 0.000883 |
| M00001 | M00909 | 0.67 | 3.44E-10 | 1.02E-09 |
| M00001 | M00950 | 0.97 | 9.88E-45 | 3.60E-43 |
| M00004 | M00010 | 0.77 | 1.43E-14 | 6.61E-14 |
| M00004 | M00015 | 0.65 | 1.57E-09 | 4.35E-09 |
| M00004 | M00018 | 0.61 | 2.30E-08 | 5.80E-08 |
| M00004 | M00019 | 0.40 | 0.000604 | 0.000921 |
| M00004 | M00021 | 0.37 | 0.001551 | 0.002235 |
| M00004 | M00023 | 0.44 | 0.000141 | 0.000237 |
| M00004 | M00024 | 0.81 | 6.47E-17 | 3.90E-16 |
| M00004 | M00028 | 0.34 | 0.004355 | 0.005857 |
| M00004 | M00048 | 0.32 | 0.007674 | 0.010008 |
| M00004 | M00049 | 0.35 | 0.003116 | 0.004278 |
| M00004 | M00050 | 0.39 | 0.000999 | 0.001474 |
| M00004 | M00052 | 0.30 | 0.012648 | 0.016036 |
| M00004 | M00083 | 0.66 | 9.72E-10 | 2.77E-09 |
| M00004 | M00086 | 0.27 | 0.022491 | 0.027418 |
| M00004 | M00096 | 0.87 | 3.09E-22 | 2.72E-21 |
| M00004 | M00097 | 0.45 | 0.000106 | 0.000182 |
| M00004 | M00112 | 0.30 | 0.012331 | 0.015647 |
| M00004 | M00115 | 0.78 | 2.65E-15 | 1.34E-14 |

| Module1 | Module2 | R | P | Padj |
|---|---|---|---|---|
| M00024 | M00165 | 0.80 | 2.07E-16 | 1.17E-15 |
| M00024 | M00175 | 0.42 | 0.000291 | 0.000467 |
| M00024 | M00176 | 0.65 | 1.15E-09 | 3.25E-09 |
| M00024 | M00307 | 0.61 | 3.15E-08 | 7.83E-08 |
| M00024 | M00364 | 0.25 | 0.035138 | 0.042028 |
| M00024 | M00432 | 0.90 | 1.76E-25 | 2.17E-24 |
| M00024 | M00527 | 0.68 | 1.99E-10 | 6.08E-10 |
| M00024 | M00549 | 0.38 | 0.001444 | 0.002089 |
| M00024 | M00621 | 0.62 | 1.05E-08 | 2.72E-08 |
| M00024 | M00793 | 0.64 | 3.71E-09 | 9.99E-09 |
| M00024 | M00842 | 0.71 | 5.72E-12 | 2.03E-11 |
| M00024 | M00843 | 0.72 | 3.92E-12 | 1.40E-11 |
| M00024 | M00844 | 0.80 | 2.38E-16 | 1.34E-15 |
| M00024 | M00854 | 0.73 | 7.59E-13 | 2.94E-12 |
| M00024 | M00880 | 0.57 | 3.09E-07 | 7.00E-07 |
| M00024 | M00881 | 0.67 | 2.26E-10 | 6.87E-10 |
| M00024 | M00909 | 0.84 | 3.73E-19 | 2.73E-18 |
| M00024 | M00932 | 0.26 | 0.028353 | 0.034334 |
| M00024 | M00950 | 0.71 | 7.35E-12 | 2.57E-11 |
| M00024 | M00970 | 0.25 | 0.040052 | 0.047593 |
| M00028 | M00048 | 0.71 | 5.16E-12 | 1.84E-11 |
| M00028 | M00049 | 0.85 | 2.50E-20 | 2.01E-19 |
| M00028 | M00050 | 0.73 | 8.64E-13 | 3.32E-12 |
| M00028 | M00052 | 0.84 | 1.54E-19 | 1.16E-18 |
| M00028 | M00083 | 0.68 | 8.91E-11 | 2.81E-10 |
| M00028 | M00086 | 0.72 | 3.68E-12 | 1.33E-11 |
| M00028 | M00096 | 0.49 | 2.07E-05 | 3.88E-05 |
| M00028 | M00097 | 0.77 | 7.42E-15 | 3.59E-14 |
| M00028 | M00112 | 0.80 | 7.70E-17 | 4.56E-16 |
| M00028 | M00115 | 0.59 | 1.18E-07 | 2.77E-07 |
| M00028 | M00118 | 0.63 | 4.84E-09 | 1.29E-08 |
| M00028 | M00119 | 0.67 | 2.32E-10 | 7.03E-10 |
| M00028 | M00120 | 0.52 | 4.04E-06 | 8.27E-06 |
| M00028 | M00121 | 0.77 | 1.20E-14 | 5.64E-14 |
| M00028 | M00133 | 0.59 | 9.86E-08 | 2.32E-07 |
| M00028 | M00135 | 0.60 | 6.22E-08 | 1.50E-07 |
| M00028 | M00145 | 0.43 | 0.000223 | 0.000364 |
| M00028 | M00155 | 0.66 | 9.82E-10 | 2.80E-09 |
| M00028 | M00157 | 0.47 | 4.12E-05 | 7.43E-05 |
| M00028 | M00161 | 0.57 | 3.34E-07 | 7.55E-07 |
| M00028 | M00165 | 0.42 | 0.000342 | 0.000543 |
| M00028 | M00175 | 0.49 | 1.88E-05 | 3.54E-05 |
| M00028 | M00176 | 0.79 | 7.81E-16 | 4.14E-15 |
| M00028 | M00307 | 0.64 | 2.41E-09 | 6.60E-09 |
| M00028 | M00364 | 0.69 | 3.69E-11 | 1.20E-10 |
| M00028 | M00432 | 0.62 | 1.71E-08 | 4.37E-08 |
| M00028 | M00527 | 0.71 | 5.95E-12 | 2.10E-11 |
| M00028 | M00549 | 0.59 | 8.30E-08 | 1.96E-07 |
| M00028 | M00621 | 0.63 | 4.80E-09 | 1.28E-08 |
| M00028 | M00793 | 0.56 | 6.86E-07 | 1.51E-06 |
| M00028 | M00842 | 0.60 | 5.63E-08 | 1.37E-07 |
| M00028 | M00843 | 0.62 | 1.48E-08 | 3.82E-08 |
| M00028 | M00844 | 0.65 | 1.36E-09 | 3.81E-09 |
| M00028 | M00854 | 0.32 | 0.007545 | 0.009848 |
| M00028 | M00880 | 0.50 | 9.65E-06 | 1.90E-05 |
| M00028 | M00881 | 0.74 | 6.44E-13 | 2.50E-12 |
| M00028 | M00909 | 0.71 | 5.74E-12 | 2.03E-11 |
| M00028 | M00932 | 0.65 | 1.46E-09 | 4.06E-09 |
| M00028 | M00950 | 0.29 | 0.01607 | 0.020042 |
| M00028 | M00970 | 0.44 | 0.00018 | 0.0003 |
| M00048 | M00049 | 0.82 | 5.58E-18 | 3.83E-17 |
| M00048 | M00050 | 0.66 | 5.19E-10 | 1.52E-09 |
| M00048 | M00052 | 0.66 | 8.11E-10 | 2.34E-09 |

| Module1 | Module2 | R | P | Padj |
|---|---|---|---|---|
| M00115 | M00970 | 0.30 | 0.011763 | 0.01504 |
| M00118 | M00119 | 0.48 | 2.64E-05 | 4.89E-05 |
| M00118 | M00120 | 0.42 | 0.000378 | 0.0006 |
| M00118 | M00121 | 0.56 | 4.62E-07 | 1.03E-06 |
| M00118 | M00133 | 0.43 | 0.000235 | 0.00038 |
| M00118 | M00135 | 0.43 | 0.000216 | 0.00035 |
| M00118 | M00145 | 0.48 | 3.58E-05 | 6.50E-05 |
| M00118 | M00155 | 0.50 | 1.25E-05 | 2.41E-05 |
| M00118 | M00157 | 0.34 | 0.003967 | 0.00535 |
| M00118 | M00161 | 0.52 | 4.13E-06 | 8.43E-06 |
| M00118 | M00165 | 0.39 | 0.001037 | 0.00153 |
| M00118 | M00175 | 0.42 | 0.000285 | 0.00046 |
| M00118 | M00176 | 0.65 | 1.32E-09 | 3.70E-09 |
| M00118 | M00307 | 0.46 | 8.44E-05 | 0.00015 |
| M00118 | M00364 | 0.56 | 4.93E-07 | 1.10E-06 |
| M00118 | M00432 | 0.38 | 0.00109 | 0.0016 |
| M00118 | M00527 | 0.51 | 6.19E-06 | 1.24E-05 |
| M00118 | M00549 | 0.52 | 3.97E-06 | 8.15E-06 |
| M00118 | M00621 | 0.47 | 4.11E-05 | 7.41E-05 |
| M00118 | M00793 | 0.37 | 0.001706 | 0.00243 |
| M00118 | M00842 | 0.44 | 0.000162 | 0.00027 |
| M00118 | M00843 | 0.46 | 8.37E-05 | 0.00015 |
| M00118 | M00844 | 0.46 | 7.14E-05 | 0.00013 |
| M00118 | M00880 | 0.52 | 5.16E-06 | 1.04E-05 |
| M00118 | M00881 | 0.55 | 1.20E-06 | 2.60E-06 |
| M00118 | M00909 | 0.49 | 1.83E-05 | 3.45E-05 |
| M00118 | M00932 | 0.64 | 3.58E-09 | 9.68E-09 |
| M00118 | M00970 | 0.37 | 0.001971 | 0.00277 |
| M00119 | M00120 | 0.47 | 5.13E-05 | 9.17E-05 |
| M00119 | M00121 | 0.93 | 7.74E-32 | 1.47E-30 |
| M00119 | M00133 | 0.60 | 4.16E-08 | 1.02E-07 |
| M00119 | M00135 | 0.98 | 1.07E-47 | 4.33E-46 |
| M00119 | M00145 | 0.40 | 0.000604 | 0.00092 |
| M00119 | M00155 | 0.97 | 2.29E-43 | 8.16E-42 |
| M00119 | M00157 | 0.28 | 0.018358 | 0.02266 |
| M00119 | M00161 | 0.32 | 0.007954 | 0.01035 |
| M00119 | M00165 | 0.44 | 0.000185 | 0.00031 |
| M00119 | M00176 | 0.83 | 1.27E-18 | 9.10E-18 |
| M00119 | M00307 | 0.37 | 0.001727 | 0.00246 |
| M00119 | M00364 | 0.44 | 0.00015 | 0.00025 |
| M00119 | M00432 | 0.75 | 9.67E-14 | 4.12E-13 |
| M00119 | M00527 | 0.99 | 1.26E-61 | 5.93E-60 |
| M00119 | M00549 | 0.33 | 0.005819 | 0.00771 |
| M00119 | M00621 | 0.71 | 1.06E-11 | 3.68E-11 |
| M00119 | M00793 | 0.98 | 1.00E-46 | 3.94E-45 |
| M00119 | M00842 | 0.80 | 8.86E-17 | 5.23E-16 |
| M00119 | M00843 | 0.82 | 1.27E-17 | 8.37E-17 |
| M00119 | M00844 | 0.75 | 1.15E-13 | 4.88E-13 |
| M00119 | M00854 | 0.29 | 0.01652 | 0.02052 |
| M00119 | M00880 | 0.79 | 4.45E-16 | 2.43E-15 |
| M00119 | M00881 | 0.89 | 1.14E-24 | 1.27E-23 |
| M00119 | M00909 | 0.88 | 1.08E-23 | 1.09E-22 |
| M00119 | M00932 | 0.46 | 6.22E-05 | 0.00011 |
| M00120 | M00121 | 0.68 | 1.17E-10 | 3.63E-10 |
| M00120 | M00133 | 0.93 | 2.19E-31 | 4.04E-30 |
| M00120 | M00135 | 0.41 | 0.000522 | 0.00081 |
| M00120 | M00145 | 0.81 | 1.93E-17 | 1.24E-16 |
| M00120 | M00155 | 0.43 | 0.000245 | 0.0004 |
| M00120 | M00157 | 0.53 | 2.69E-06 | 5.66E-06 |
| M00120 | M00161 | 0.38 | 0.001298 | 0.00189 |
| M00120 | M00163 | 0.81 | 7.61E-17 | 4.53E-16 |
| M00120 | M00165 | 0.95 | 1.99E-36 | 5.08E-35 |
| M00120 | M00175 | 0.46 | 6.69E-05 | 0.00012 |

| Module1 | Module2 | R | P | Padj |
|---|---|---|---|---|
| M00004 | M00119 | 0.39 | 0.000991 | 0.001464 |
| M00004 | M00120 | 0.89 | 6.81E-25 | 7.64E-24 |
| M00004 | M00121 | 0.56 | 5.11E-07 | 1.13E-06 |
| M00004 | M00133 | 0.91 | 7.11E-28 | 1.06E-26 |
| M00004 | M00135 | 0.39 | 0.000984 | 0.001454 |
| M00004 | M00145 | 0.78 | 5.01E-15 | 2.47E-14 |
| M00004 | M00155 | 0.40 | 0.000627 | 0.000953 |
| M00004 | M00157 | 0.58 | 2.04E-07 | 4.71E-07 |
| M00004 | M00163 | 0.92 | 2.46E-28 | 3.77E-27 |
| M00004 | M00165 | 0.94 | 5.83E-32 | 1.12E-30 |
| M00004 | M00175 | 0.40 | 0.000577 | 0.000883 |
| M00004 | M00176 | 0.48 | 3.01E-05 | 5.57E-05 |
| M00004 | M00307 | 0.41 | 0.000449 | 0.000706 |
| M00004 | M00432 | 0.88 | 6.19E-23 | 5.79E-22 |
| M00004 | M00527 | 0.38 | 0.001226 | 0.001791 |
| M00004 | M00621 | 0.30 | 0.013315 | 0.016808 |
| M00004 | M00793 | 0.35 | 0.003602 | 0.004895 |
| M00004 | M00842 | 0.40 | 0.000735 | 0.001104 |
| M00004 | M00843 | 0.40 | 0.000712 | 0.001071 |
| M00004 | M00844 | 0.75 | 1.54E-13 | 6.45E-13 |
| M00004 | M00854 | 0.96 | 1.62E-37 | 4.29E-36 |
| M00004 | M00880 | 0.38 | 0.00116 | 0.001698 |
| M00004 | M00881 | 0.40 | 0.000611 | 0.000931 |
| M00004 | M00909 | 0.66 | 5.04E-10 | 1.48E-09 |
| M00004 | M00950 | 0.96 | 2.47E-37 | 6.43E-36 |
| M00010 | M00015 | 0.68 | 1.14E-10 | 3.57E-10 |
| M00010 | M00018 | 0.60 | 5.88E-08 | 1.43E-07 |
| M00010 | M00019 | 0.41 | 0.000522 | 0.000807 |
| M00010 | M00021 | 0.53 | 2.45E-06 | 5.19E-06 |
| M00010 | M00022 | 0.37 | 0.001803 | 0.002559 |
| M00010 | M00023 | 0.66 | 8.41E-10 | 2.41E-09 |
| M00010 | M00024 | 0.81 | 3.05E-17 | 1.90E-16 |
| M00010 | M00028 | 0.56 | 5.49E-07 | 1.21E-06 |
| M00010 | M00048 | 0.47 | 3.84E-05 | 6.94E-05 |
| M00010 | M00049 | 0.48 | 3.41E-05 | 6.21E-05 |
| M00010 | M00050 | 0.50 | 1.38E-05 | 2.66E-05 |
| M00010 | M00052 | 0.51 | 7.60E-06 | 1.51E-05 |
| M00010 | M00083 | 0.62 | 9.57E-09 | 2.51E-08 |
| M00010 | M00086 | 0.39 | 0.000802 | 0.001198 |
| M00010 | M00096 | 0.72 | 2.61E-12 | 9.61E-12 |
| M00010 | M00097 | 0.57 | 2.44E-07 | 5.60E-07 |
| M00010 | M00112 | 0.44 | 0.000135 | 0.000228 |
| M00010 | M00115 | 0.77 | 1.40E-14 | 6.52E-14 |
| M00010 | M00118 | 0.47 | 3.74E-05 | 6.78E-05 |
| M00010 | M00119 | 0.42 | 0.000362 | 0.000573 |
| M00010 | M00120 | 0.78 | 3.75E-15 | 1.87E-14 |
| M00010 | M00121 | 0.57 | 3.50E-07 | 7.91E-07 |
| M00010 | M00133 | 0.80 | 1.85E-16 | 1.06E-15 |
| M00010 | M00135 | 0.44 | 0.000141 | 0.000237 |
| M00010 | M00145 | 0.76 | 4.64E-14 | 2.03E-13 |
| M00010 | M00155 | 0.43 | 0.000261 | 0.000421 |
| M00010 | M00157 | 0.82 | 5.21E-18 | 3.62E-17 |
| M00010 | M00161 | 0.38 | 0.001245 | 0.001816 |
| M00010 | M00163 | 0.64 | 3.47E-09 | 9.43E-09 |
| M00010 | M00165 | 0.79 | 5.50E-16 | 2.98E-15 |
| M00010 | M00175 | 0.44 | 0.000167 | 0.00028 |
| M00010 | M00176 | 0.55 | 9.33E-07 | 2.04E-06 |
| M00010 | M00307 | 0.66 | 5.19E-10 | 1.52E-09 |
| M00010 | M00364 | 0.28 | 0.019154 | 0.023527 |
| M00010 | M00432 | 0.76 | 2.12E-14 | 9.70E-14 |
| M00010 | M00527 | 0.45 | 9.70E-05 | 0.000167 |
| M00010 | M00549 | 0.40 | 0.000627 | 0.000953 |
| M00010 | M00621 | 0.50 | 1.04E-05 | 2.04E-05 |

| Module1 | Module2 | R | P | Padj |
|---|---|---|---|---|
| M00048 | M00083 | 0.62 | 9.86E-09 | 2.58E-08 |
| M00048 | M00086 | 0.60 | 6.58E-08 | 1.58E-07 |
| M00048 | M00096 | 0.48 | 3.11E-05 | 5.72E-05 |
| M00048 | M00097 | 0.62 | 1.46E-08 | 3.77E-08 |
| M00048 | M00112 | 0.76 | 5.73E-14 | 2.49E-13 |
| M00048 | M00115 | 0.50 | 1.31E-05 | 2.53E-05 |
| M00048 | M00118 | 0.73 | 7.99E-13 | 3.08E-12 |
| M00048 | M00119 | 0.62 | 1.16E-08 | 3.01E-08 |
| M00048 | M00120 | 0.62 | 1.76E-08 | 4.47E-08 |
| M00048 | M00121 | 0.73 | 1.01E-12 | 3.85E-12 |
| M00048 | M00133 | 0.56 | 5.59E-07 | 1.24E-06 |
| M00048 | M00135 | 0.52 | 3.76E-06 | 7.78E-06 |
| M00048 | M00145 | 0.61 | 1.93E-08 | 4.91E-08 |
| M00048 | M00155 | 0.58 | 2.10E-07 | 4.86E-07 |
| M00048 | M00161 | 0.46 | 6.76E-05 | 0.00012 |
| M00048 | M00165 | 0.54 | 1.68E-06 | 3.58E-06 |
| M00048 | M00175 | 0.41 | 0.000476 | 0.000743 |
| M00048 | M00176 | 0.87 | 1.39E-22 | 1.25E-21 |
| M00048 | M00307 | 0.40 | 0.000563 | 0.000865 |
| M00048 | M00364 | 0.79 | 3.47E-16 | 1.91E-15 |
| M00048 | M00432 | 0.53 | 2.51E-06 | 5.30E-06 |
| M00048 | M00527 | 0.65 | 1.06E-09 | 3.00E-09 |
| M00048 | M00549 | 0.43 | 0.000232 | 0.000378 |
| M00048 | M00621 | 0.43 | 0.000211 | 0.000346 |
| M00048 | M00793 | 0.48 | 3.24E-05 | 5.94E-05 |
| M00048 | M00842 | 0.40 | 0.000701 | 0.001055 |
| M00048 | M00843 | 0.43 | 0.00027 | 0.000434 |
| M00048 | M00844 | 0.53 | 2.77E-06 | 5.81E-06 |
| M00048 | M00854 | 0.30 | 0.012172 | 0.015486 |
| M00048 | M00880 | 0.40 | 0.000745 | 0.001118 |
| M00048 | M00881 | 0.65 | 1.55E-09 | 4.31E-09 |
| M00048 | M00909 | 0.59 | 7.84E-08 | 1.86E-07 |
| M00048 | M00932 | 0.90 | 2.05E-25 | 2.49E-24 |
| M00048 | M00950 | 0.32 | 0.007043 | 0.009235 |
| M00048 | M00970 | 0.28 | 0.021721 | 0.026501 |
| M00049 | M00050 | 0.83 | 1.57E-18 | 1.11E-17 |
| M00049 | M00052 | 0.83 | 5.54E-19 | 4.02E-18 |
| M00049 | M00083 | 0.83 | 1.60E-18 | 1.14E-17 |
| M00049 | M00086 | 0.65 | 2.14E-09 | 5.88E-09 |
| M00049 | M00096 | 0.49 | 1.80E-05 | 3.40E-05 |
| M00049 | M00097 | 0.87 | 3.26E-22 | 2.85E-21 |
| M00049 | M00112 | 0.89 | 2.33E-24 | 2.51E-23 |
| M00049 | M00115 | 0.48 | 2.85E-05 | 5.28E-05 |
| M00049 | M00118 | 0.65 | 1.21E-09 | 3.39E-09 |
| M00049 | M00119 | 0.85 | 2.86E-20 | 2.26E-19 |
| M00049 | M00120 | 0.55 | 7.64E-07 | 1.68E-06 |
| M00049 | M00121 | 0.91 | 7.67E-28 | 1.12E-26 |
| M00049 | M00133 | 0.58 | 1.52E-07 | 3.53E-07 |
| M00049 | M00135 | 0.76 | 3.84E-14 | 1.72E-13 |
| M00049 | M00145 | 0.40 | 0.000587 | 0.000897 |
| M00049 | M00155 | 0.82 | 9.40E-18 | 6.26E-17 |
| M00049 | M00157 | 0.33 | 0.005537 | 0.007359 |
| M00049 | M00161 | 0.56 | 4.75E-07 | 1.06E-06 |
| M00049 | M00165 | 0.45 | 0.000112 | 0.000191 |
| M00049 | M00175 | 0.45 | 0.000121 | 0.000205 |
| M00049 | M00176 | 0.90 | 1.74E-25 | 2.17E-24 |
| M00049 | M00307 | 0.52 | 4.16E-06 | 8.48E-06 |
| M00049 | M00364 | 0.77 | 5.88E-15 | 2.90E-14 |
| M00049 | M00432 | 0.66 | 7.48E-10 | 2.16E-09 |
| M00049 | M00527 | 0.89 | 5.25E-24 | 5.51E-23 |
| M00049 | M00549 | 0.53 | 2.85E-06 | 5.98E-06 |
| M00049 | M00621 | 0.68 | 1.14E-10 | 3.56E-10 |
| M00049 | M00793 | 0.74 | 3.21E-13 | 1.29E-12 |

| Module1 | Module2 | R | P | Padj |
|---|---|---|---|---|
| M00120 | M00176 | 0.68 | 1.05E-10 | 3.30E-10 |
| M00120 | M00307 | 0.49 | 1.96E-05 | 3.67E-05 |
| M00120 | M00364 | 0.53 | 2.88E-06 | 6.02E-06 |
| M00120 | M00432 | 0.86 | 5.93E-21 | 4.99E-20 |
| M00120 | M00527 | 0.48 | 2.43E-05 | 4.53E-05 |
| M00120 | M00549 | 0.26 | 0.034371 | 0.04114 |
| M00120 | M00621 | 0.33 | 0.005748 | 0.00763 |
| M00120 | M00793 | 0.35 | 0.002823 | 0.00389 |
| M00120 | M00842 | 0.43 | 0.000204 | 0.00034 |
| M00120 | M00843 | 0.44 | 0.000139 | 0.00023 |
| M00120 | M00844 | 0.72 | 3.63E-12 | 1.31E-11 |
| M00120 | M00854 | 0.87 | 9.18E-23 | 8.31E-22 |
| M00120 | M00880 | 0.36 | 0.002635 | 0.00365 |
| M00120 | M00881 | 0.51 | 7.52E-06 | 1.50E-05 |
| M00120 | M00909 | 0.65 | 1.39E-09 | 3.87E-09 |
| M00120 | M00932 | 0.54 | 1.57E-06 | 3.37E-06 |
| M00120 | M00950 | 0.90 | 1.46E-25 | 1.85E-24 |
| M00121 | M00133 | 0.75 | 9.06E-14 | 3.87E-13 |
| M00121 | M00135 | 0.88 | 2.00E-23 | 1.93E-22 |
| M00121 | M00145 | 0.53 | 2.68E-06 | 5.65E-06 |
| M00121 | M00155 | 0.90 | 8.56E-26 | 1.10E-24 |
| M00121 | M00157 | 0.41 | 0.000528 | 0.00081 |
| M00121 | M00161 | 0.43 | 0.000262 | 0.00042 |
| M00121 | M00163 | 0.25 | 0.035898 | 0.0429 |
| M00121 | M00165 | 0.63 | 6.81E-09 | 1.81E-08 |
| M00121 | M00175 | 0.34 | 0.003766 | 0.00509 |
| M00121 | M00176 | 0.91 | 3.91E-27 | 5.43E-26 |
| M00121 | M00307 | 0.52 | 5.82E-06 | 1.17E-05 |
| M00121 | M00364 | 0.59 | 8.24E-08 | 1.95E-07 |
| M00121 | M00432 | 0.84 | 1.55E-19 | 1.16E-18 |
| M00121 | M00527 | 0.94 | 1.03E-33 | 2.13E-32 |
| M00121 | M00549 | 0.38 | 0.001479 | 0.00214 |
| M00121 | M00621 | 0.71 | 1.00E-11 | 3.49E-11 |
| M00121 | M00793 | 0.87 | 7.57E-22 | 6.53E-21 |
| M00121 | M00842 | 0.74 | 3.68E-13 | 1.47E-12 |
| M00121 | M00843 | 0.76 | 4.50E-14 | 1.98E-13 |
| M00121 | M00844 | 0.82 | 9.31E-18 | 6.22E-17 |
| M00121 | M00854 | 0.50 | 1.28E-05 | 2.47E-05 |
| M00121 | M00880 | 0.76 | 4.46E-14 | 1.97E-13 |
| M00121 | M00881 | 0.91 | 1.07E-27 | 1.52E-26 |
| M00121 | M00909 | 0.89 | 5.84E-25 | 6.61E-24 |
| M00121 | M00932 | 0.61 | 1.97E-08 | 4.99E-08 |
| M00121 | M00950 | 0.45 | 9.72E-05 | 0.00017 |
| M00133 | M00135 | 0.57 | 3.93E-07 | 8.85E-07 |
| M00133 | M00145 | 0.84 | 1.44E-19 | 1.10E-18 |
| M00133 | M00155 | 0.60 | 6.67E-08 | 1.60E-07 |
| M00133 | M00157 | 0.61 | 2.99E-08 | 7.45E-08 |
| M00133 | M00161 | 0.39 | 0.000846 | 0.00126 |
| M00133 | M00163 | 0.75 | 7.14E-14 | 3.06E-13 |
| M00133 | M00165 | 0.94 | 4.71E-34 | 1.02E-32 |
| M00133 | M00175 | 0.51 | 7.13E-06 | 1.42E-05 |
| M00133 | M00176 | 0.70 | 3.00E-11 | 9.84E-11 |
| M00133 | M00307 | 0.53 | 3.19E-06 | 6.64E-06 |
| M00133 | M00364 | 0.41 | 0.000513 | 0.0008 |
| M00133 | M00432 | 0.94 | 3.36E-34 | 7.54E-33 |
| M00133 | M00527 | 0.60 | 6.00E-08 | 1.45E-07 |
| M00133 | M00549 | 0.39 | 0.001076 | 0.00158 |
| M00133 | M00621 | 0.47 | 4.20E-05 | 7.56E-05 |
| M00133 | M00793 | 0.52 | 5.51E-06 | 1.11E-05 |
| M00133 | M00842 | 0.57 | 2.73E-07 | 6.23E-07 |
| M00133 | M00843 | 0.57 | 2.59E-07 | 5.93E-07 |
| M00133 | M00844 | 0.84 | 2.38E-19 | 1.77E-18 |
| M00133 | M00854 | 0.89 | 3.59E-24 | 3.82E-23 |

| Module1 | Module2 | R | P | Padj |
|---|---|---|---|---|
| M00010 | M00793 | 0.35 | 0.003392 | 0.004622 |
| M00010 | M00842 | 0.60 | 5.84E-08 | 1.42E-07 |
| M00010 | M00843 | 0.62 | 1.69E-08 | 4.32E-08 |
| M00010 | M00844 | 0.63 | 5.01E-09 | 1.33E-08 |
| M00010 | M00854 | 0.74 | 4.22E-13 | 1.66E-12 |
| M00010 | M00880 | 0.38 | 0.001477 | 0.002135 |
| M00010 | M00881 | 0.48 | 2.45E-05 | 4.57E-05 |
| M00010 | M00909 | 0.65 | 1.09E-09 | 3.07E-09 |
| M00010 | M00932 | 0.30 | 0.013214 | 0.016695 |
| M00010 | M00950 | 0.75 | 1.75E-13 | 7.30E-13 |
| M00010 | M00970 | 0.31 | 0.010211 | 0.013182 |
| M00015 | M00018 | 0.98 | 5.07E-48 | 2.11E-46 |
| M00015 | M00019 | 0.91 | 8.61E-28 | 1.24E-26 |
| M00015 | M00021 | 0.77 | 7.22E-15 | 3.52E-14 |
| M00015 | M00023 | 0.67 | 4.33E-10 | 1.28E-09 |
| M00015 | M00024 | 0.84 | 1.08E-19 | 8.27E-19 |
| M00015 | M00028 | 0.63 | 8.03E-09 | 2.11E-08 |
| M00015 | M00048 | 0.49 | 1.64E-05 | 3.12E-05 |
| M00015 | M00049 | 0.73 | 1.36E-12 | 5.09E-12 |
| M00015 | M00050 | 0.88 | 8.06E-23 | 7.43E-22 |
| M00015 | M00052 | 0.72 | 2.94E-12 | 1.07E-11 |
| M00015 | M00083 | 0.95 | 1.86E-35 | 4.45E-34 |
| M00015 | M00086 | 0.35 | 0.003581 | 0.004871 |
| M00015 | M00096 | 0.62 | 1.59E-08 | 4.06E-08 |
| M00015 | M00097 | 0.88 | 6.33E-24 | 6.50E-23 |
| M00015 | M00112 | 0.72 | 3.41E-12 | 1.24E-11 |
| M00015 | M00115 | 0.52 | 4.97E-06 | 1.01E-05 |
| M00015 | M00118 | 0.44 | 0.000176 | 0.000294 |
| M00015 | M00119 | 0.89 | 5.85E-25 | 6.61E-24 |
| M00015 | M00120 | 0.62 | 1.07E-08 | 2.77E-08 |
| M00015 | M00121 | 0.89 | 6.05E-24 | 6.26E-23 |
| M00015 | M00133 | 0.76 | 2.46E-14 | 1.12E-13 |
| M00015 | M00135 | 0.92 | 9.10E-29 | 1.47E-27 |
| M00015 | M00145 | 0.54 | 1.68E-06 | 3.58E-06 |
| M00015 | M00155 | 0.88 | 1.81E-23 | 1.77E-22 |
| M00015 | M00157 | 0.52 | 4.38E-06 | 8.91E-06 |
| M00015 | M00161 | 0.32 | 0.007459 | 0.009745 |
| M00015 | M00163 | 0.35 | 0.003354 | 0.004579 |
| M00015 | M00165 | 0.62 | 1.13E-08 | 2.93E-08 |
| M00015 | M00175 | 0.28 | 0.018418 | 0.022719 |
| M00015 | M00176 | 0.77 | 1.04E-14 | 4.94E-14 |
| M00015 | M00307 | 0.49 | 1.63E-05 | 3.11E-05 |
| M00015 | M00364 | 0.29 | 0.015482 | 0.019376 |
| M00015 | M00432 | 0.89 | 5.94E-24 | 6.18E-23 |
| M00015 | M00527 | 0.90 | 2.00E-25 | 2.45E-24 |
| M00015 | M00549 | 0.35 | 0.00368 | 0.004991 |
| M00015 | M00621 | 0.70 | 2.55E-11 | 8.52E-11 |
| M00015 | M00793 | 0.89 | 2.48E-24 | 2.65E-23 |
| M00015 | M00842 | 0.82 | 3.50E-18 | 2.46E-17 |
| M00015 | M00843 | 0.85 | 2.61E-20 | 2.08E-19 |
| M00015 | M00844 | 0.82 | 6.99E-18 | 4.74E-17 |
| M00015 | M00854 | 0.54 | 1.30E-06 | 2.80E-06 |
| M00015 | M00880 | 0.82 | 3.83E-18 | 2.67E-17 |
| M00015 | M00881 | 0.81 | 2.53E-17 | 1.61E-16 |
| M00015 | M00909 | 0.92 | 1.48E-28 | 2.34E-27 |
| M00015 | M00932 | 0.30 | 0.011132 | 0.014313 |
| M00015 | M00950 | 0.49 | 1.60E-05 | 3.05E-05 |
| M00018 | M00019 | 0.96 | 8.26E-40 | 2.56E-38 |
| M00018 | M00021 | 0.79 | 4.23E-16 | 2.32E-15 |
| M00018 | M00023 | 0.62 | 1.05E-08 | 2.72E-08 |
| M00018 | M00024 | 0.82 | 1.01E-17 | 6.69E-17 |
| M00018 | M00028 | 0.61 | 2.80E-08 | 7.02E-08 |
| M00018 | M00048 | 0.52 | 3.96E-06 | 8.14E-06 |

| Module1 | Module2 | R | P | Padj |
|---|---|---|---|---|
| M00049 | M00842 | 0.64 | 2.64E-09 | 7.22E-09 |
| M00049 | M00843 | 0.67 | 3.25E-10 | 9.73E-10 |
| M00049 | M00844 | 0.72 | 4.50E-12 | 1.61E-11 |
| M00049 | M00854 | 0.30 | 0.012112 | 0.015422 |
| M00049 | M00880 | 0.64 | 3.62E-09 | 9.78E-09 |
| M00049 | M00881 | 0.85 | 2.56E-20 | 2.05E-19 |
| M00049 | M00909 | 0.80 | 1.33E-16 | 7.61E-16 |
| M00049 | M00932 | 0.74 | 2.70E-13 | 1.10E-12 |
| M00049 | M00950 | 0.26 | 0.029963 | 0.036164 |
| M00049 | M00970 | 0.30 | 0.01307 | 0.016542 |
| M00050 | M00052 | 0.86 | 3.27E-21 | 2.79E-20 |
| M00050 | M00083 | 0.89 | 1.58E-24 | 1.73E-23 |
| M00050 | M00086 | 0.46 | 6.77E-05 | 0.00012 |
| M00050 | M00096 | 0.43 | 0.000206 | 0.000338 |
| M00050 | M00097 | 0.92 | 2.09E-28 | 3.24E-27 |
| M00050 | M00112 | 0.78 | 1.62E-15 | 8.35E-15 |
| M00050 | M00115 | 0.37 | 0.001939 | 0.002732 |
| M00050 | M00118 | 0.56 | 4.82E-07 | 1.07E-06 |
| M00050 | M00119 | 0.95 | 1.36E-34 | 3.16E-33 |
| M00050 | M00120 | 0.45 | 9.81E-05 | 0.000168 |
| M00050 | M00121 | 0.89 | 1.60E-24 | 1.74E-23 |
| M00050 | M00133 | 0.61 | 2.68E-08 | 6.72E-08 |
| M00050 | M00135 | 0.94 | 8.86E-34 | 1.87E-32 |
| M00050 | M00145 | 0.48 | 3.36E-05 | 6.13E-05 |
| M00050 | M00155 | 0.96 | 2.49E-39 | 7.26E-38 |
| M00050 | M00157 | 0.38 | 0.001399 | 0.002029 |
| M00050 | M00161 | 0.42 | 0.000336 | 0.000537 |
| M00050 | M00165 | 0.45 | 8.63E-05 | 0.000149 |
| M00050 | M00175 | 0.30 | 0.013729 | 0.017315 |
| M00050 | M00176 | 0.85 | 5.73E-20 | 4.42E-19 |
| M00050 | M00307 | 0.48 | 3.09E-05 | 5.69E-05 |
| M00050 | M00364 | 0.41 | 0.000542 | 0.000833 |
| M00050 | M00432 | 0.73 | 7.60E-13 | 2.94E-12 |
| M00050 | M00527 | 0.94 | 7.80E-34 | 1.67E-32 |
| M00050 | M00549 | 0.40 | 0.000665 | 0.001005 |
| M00050 | M00621 | 0.74 | 2.34E-13 | 9.60E-13 |
| M00050 | M00793 | 0.92 | 1.80E-28 | 2.83E-27 |
| M00050 | M00842 | 0.80 | 9.75E-17 | 5.73E-16 |
| M00050 | M00843 | 0.82 | 5.65E-18 | 3.85E-17 |
| M00050 | M00844 | 0.76 | 5.94E-14 | 2.58E-13 |
| M00050 | M00854 | 0.28 | 0.020151 | 0.024669 |
| M00050 | M00880 | 0.75 | 1.17E-13 | 4.94E-13 |
| M00050 | M00881 | 0.88 | 2.20E-23 | 2.10E-22 |
| M00050 | M00909 | 0.88 | 5.59E-23 | 5.26E-22 |
| M00050 | M00932 | 0.48 | 3.09E-05 | 5.69E-05 |
| M00052 | M00083 | 0.73 | 1.16E-12 | 4.37E-12 |
| M00052 | M00086 | 0.62 | 1.21E-08 | 3.13E-08 |
| M00052 | M00096 | 0.37 | 0.001577 | 0.002261 |
| M00052 | M00097 | 0.89 | 4.52E-25 | 5.19E-24 |
| M00052 | M00112 | 0.79 | 7.68E-16 | 4.09E-15 |
| M00052 | M00115 | 0.46 | 6.78E-05 | 0.00012 |
| M00052 | M00118 | 0.66 | 6.70E-10 | 1.94E-09 |
| M00052 | M00119 | 0.77 | 8.63E-15 | 4.16E-14 |
| M00052 | M00120 | 0.39 | 0.001013 | 0.001494 |
| M00052 | M00121 | 0.77 | 1.27E-14 | 5.92E-14 |
| M00052 | M00133 | 0.54 | 1.85E-06 | 3.94E-06 |
| M00052 | M00135 | 0.74 | 4.07E-13 | 1.61E-12 |
| M00052 | M00145 | 0.41 | 0.000432 | 0.000681 |
| M00052 | M00155 | 0.81 | 1.57E-17 | 1.02E-16 |
| M00052 | M00157 | 0.46 | 8.19E-05 | 0.000143 |
| M00052 | M00161 | 0.55 | 1.27E-06 | 2.74E-06 |
| M00052 | M00165 | 0.36 | 0.002527 | 0.003508 |
| M00052 | M00175 | 0.50 | 1.52E-05 | 2.91E-05 |

| Module1 | Module2 | R | P | Padj |
|---|---|---|---|---|
| M00133 | M00880 | 0.53 | 3.60E-06 | 7.48E-06 |
| M00133 | M00881 | 0.61 | 2.27E-08 | 5.74E-08 |
| M00133 | M00909 | 0.81 | 7.08E-17 | 4.23E-16 |
| M00133 | M00932 | 0.43 | 0.000194 | 0.00032 |
| M00133 | M00950 | 0.87 | 9.03E-23 | 8.23E-22 |
| M00135 | M00145 | 0.40 | 0.000672 | 0.00101 |
| M00135 | M00155 | 0.97 | 3.57E-41 | 1.18E-39 |
| M00135 | M00157 | 0.29 | 0.014495 | 0.01823 |
| M00135 | M00161 | 0.25 | 0.036693 | 0.04382 |
| M00135 | M00165 | 0.41 | 0.000494 | 0.00077 |
| M00135 | M00176 | 0.76 | 3.55E-14 | 1.61E-13 |
| M00135 | M00307 | 0.34 | 0.003749 | 0.00508 |
| M00135 | M00364 | 0.28 | 0.019562 | 0.02399 |
| M00135 | M00432 | 0.74 | 3.95E-13 | 1.57E-12 |
| M00135 | M00527 | 0.97 | 1.46E-41 | 4.93E-40 |
| M00135 | M00549 | 0.29 | 0.016439 | 0.02043 |
| M00135 | M00621 | 0.70 | 1.66E-11 | 5.63E-11 |
| M00135 | M00793 | 0.99 | 8.95E-61 | 4.08E-59 |
| M00135 | M00842 | 0.82 | 5.59E-18 | 3.83E-17 |
| M00135 | M00843 | 0.84 | 4.34E-19 | 3.16E-18 |
| M00135 | M00844 | 0.72 | 2.23E-12 | 8.25E-12 |
| M00135 | M00854 | 0.27 | 0.0243 | 0.0295 |
| M00135 | M00880 | 0.81 | 3.77E-17 | 2.32E-16 |
| M00135 | M00881 | 0.86 | 4.97E-21 | 4.22E-20 |
| M00135 | M00909 | 0.87 | 1.53E-22 | 1.37E-21 |
| M00135 | M00932 | 0.32 | 0.007339 | 0.0096 |
| M00145 | M00155 | 0.42 | 0.000306 | 0.00049 |
| M00145 | M00157 | 0.45 | 9.33E-05 | 0.00016 |
| M00145 | M00161 | 0.31 | 0.008905 | 0.01155 |
| M00145 | M00163 | 0.69 | 6.21E-11 | 2.00E-10 |
| M00145 | M00165 | 0.90 | 1.25E-25 | 1.60E-24 |
| M00145 | M00175 | 0.52 | 3.83E-06 | 7.90E-06 |
| M00145 | M00176 | 0.55 | 1.14E-06 | 2.47E-06 |
| M00145 | M00307 | 0.37 | 0.001903 | 0.00269 |
| M00145 | M00364 | 0.26 | 0.033372 | 0.04005 |
| M00145 | M00432 | 0.76 | 4.36E-14 | 1.93E-13 |
| M00145 | M00527 | 0.39 | 0.000868 | 0.00129 |
| M00145 | M00549 | 0.43 | 0.00026 | 0.00042 |
| M00145 | M00621 | 0.33 | 0.005516 | 0.00734 |
| M00145 | M00793 | 0.33 | 0.005917 | 0.00782 |
| M00145 | M00842 | 0.37 | 0.001569 | 0.00226 |
| M00145 | M00843 | 0.37 | 0.001952 | 0.00275 |
| M00145 | M00844 | 0.67 | 2.80E-10 | 8.43E-10 |
| M00145 | M00854 | 0.78 | 1.53E-15 | 7.94E-15 |
| M00145 | M00880 | 0.34 | 0.00375 | 0.00508 |
| M00145 | M00881 | 0.46 | 8.47E-05 | 0.00015 |
| M00145 | M00909 | 0.66 | 8.24E-10 | 2.37E-09 |
| M00145 | M00932 | 0.38 | 0.00143 | 0.00207 |
| M00145 | M00950 | 0.79 | 6.60E-16 | 3.52E-15 |
| M00145 | M00970 | 0.35 | 0.003629 | 0.00493 |
| M00155 | M00157 | 0.31 | 0.009987 | 0.01291 |
| M00155 | M00161 | 0.37 | 0.001934 | 0.00273 |
| M00155 | M00165 | 0.43 | 0.000199 | 0.00033 |
| M00155 | M00175 | 0.30 | 0.011304 | 0.0145 |
| M00155 | M00176 | 0.78 | 2.34E-15 | 1.19E-14 |
| M00155 | M00307 | 0.39 | 0.000851 | 0.00126 |
| M00155 | M00364 | 0.36 | 0.002378 | 0.00331 |
| M00155 | M00432 | 0.74 | 4.23E-13 | 1.66E-12 |
| M00155 | M00527 | 0.96 | 4.00E-39 | 1.14E-37 |
| M00155 | M00549 | 0.41 | 0.000457 | 0.00071 |
| M00155 | M00621 | 0.76 | 3.69E-14 | 1.67E-13 |
| M00155 | M00793 | 0.96 | 2.92E-38 | 7.90E-37 |
| M00155 | M00842 | 0.80 | 1.01E-16 | 5.90E-16 |

| Module1 | Module2 | R | P | Padj |
|---|---|---|---|---|
| M00018 | M00049 | 0.75 | 1.91E-13 | 7.91E-13 |
| M00018 | M00050 | 0.91 | 8.60E-27 | 1.18E-25 |
| M00018 | M00052 | 0.71 | 1.07E-11 | 3.68E-11 |
| M00018 | M00083 | 0.98 | 1.18E-45 | 4.54E-44 |
| M00018 | M00086 | 0.32 | 0.006659 | 0.008763 |
| M00018 | M00096 | 0.58 | 1.32E-07 | 3.09E-07 |
| M00018 | M00097 | 0.89 | 1.30E-24 | 1.44E-23 |
| M00018 | M00112 | 0.70 | 3.25E-11 | 1.06E-10 |
| M00018 | M00115 | 0.44 | 0.000183 | 0.000304 |
| M00018 | M00118 | 0.42 | 0.000385 | 0.000609 |
| M00018 | M00119 | 0.94 | 3.78E-34 | 8.36E-33 |
| M00018 | M00120 | 0.60 | 6.90E-08 | 1.65E-07 |
| M00018 | M00121 | 0.91 | 9.66E-28 | 1.38E-26 |
| M00018 | M00133 | 0.74 | 3.88E-13 | 1.55E-12 |
| M00018 | M00135 | 0.96 | 1.53E-39 | 4.54E-38 |
| M00018 | M00145 | 0.54 | 1.83E-06 | 3.89E-06 |
| M00018 | M00155 | 0.93 | 1.51E-30 | 2.62E-29 |
| M00018 | M00157 | 0.43 | 0.000233 | 0.00038 |
| M00018 | M00161 | 0.26 | 0.029867 | 0.036108 |
| M00018 | M00163 | 0.30 | 0.011974 | 0.015274 |
| M00018 | M00165 | 0.60 | 3.99E-08 | 9.85E-08 |
| M00018 | M00175 | 0.25 | 0.039261 | 0.046767 |
| M00018 | M00176 | 0.78 | 1.97E-15 | 1.00E-14 |
| M00018 | M00307 | 0.41 | 0.000427 | 0.000673 |
| M00018 | M00364 | 0.28 | 0.017967 | 0.022219 |
| M00018 | M00432 | 0.88 | 1.33E-23 | 1.32E-22 |
| M00018 | M00527 | 0.93 | 9.45E-32 | 1.77E-30 |
| M00018 | M00549 | 0.30 | 0.011105 | 0.01429 |
| M00018 | M00621 | 0.68 | 1.27E-10 | 3.90E-10 |
| M00018 | M00793 | 0.95 | 2.25E-34 | 5.13E-33 |
| M00018 | M00842 | 0.81 | 2.42E-17 | 1.55E-16 |
| M00018 | M00843 | 0.83 | 1.99E-18 | 1.40E-17 |
| M00018 | M00844 | 0.82 | 9.08E-18 | 6.10E-17 |
| M00018 | M00854 | 0.50 | 1.01E-05 | 1.97E-05 |
| M00018 | M00880 | 0.81 | 3.00E-17 | 1.88E-16 |
| M00018 | M00881 | 0.84 | 3.34E-19 | 2.46E-18 |
| M00018 | M00909 | 0.93 | 7.66E-31 | 1.38E-29 |
| M00018 | M00932 | 0.32 | 0.0074 | 0.009677 |
| M00018 | M00950 | 0.46 | 7.42E-05 | 0.00013 |
| M00019 | M00021 | 0.81 | 6.17E-17 | 3.73E-16 |
| M00019 | M00023 | 0.56 | 4.34E-07 | 9.74E-07 |
| M00019 | M00024 | 0.69 | 4.57E-11 | 1.48E-10 |
| M00019 | M00028 | 0.59 | 9.19E-08 | 2.17E-07 |
| M00019 | M00048 | 0.50 | 1.52E-05 | 2.91E-05 |
| M00019 | M00049 | 0.76 | 4.49E-14 | 1.98E-13 |
| M00019 | M00050 | 0.93 | 9.71E-31 | 1.73E-29 |
| M00019 | M00052 | 0.73 | 1.42E-12 | 5.30E-12 |
| M00019 | M00083 | 0.93 | 1.45E-30 | 2.54E-29 |
| M00019 | M00086 | 0.30 | 0.010884 | 0.014018 |
| M00019 | M00096 | 0.41 | 0.000472 | 0.000737 |
| M00019 | M00097 | 0.90 | 2.67E-25 | 3.14E-24 |
| M00019 | M00112 | 0.70 | 3.23E-11 | 1.06E-10 |
| M00019 | M00115 | 0.25 | 0.039902 | 0.047453 |
| M00019 | M00118 | 0.39 | 0.000832 | 0.001239 |
| M00019 | M00119 | 0.98 | 2.51E-49 | 1.08E-47 |
| M00019 | M00120 | 0.41 | 0.000553 | 0.00085 |
| M00019 | M00121 | 0.89 | 4.37E-24 | 4.62E-23 |
| M00019 | M00133 | 0.57 | 3.06E-07 | 6.95E-07 |
| M00019 | M00135 | 0.99 | 3.13E-65 | 1.52E-63 |
| M00019 | M00145 | 0.38 | 0.001368 | 0.001987 |
| M00019 | M00155 | 0.97 | 1.04E-41 | 3.60E-40 |
| M00019 | M00157 | 0.28 | 0.019075 | 0.023449 |
| M00019 | M00165 | 0.41 | 0.000524 | 0.00081 |
| M00052 | M00176 | 0.76 | 6.18E-14 | 2.67E-13 |
| M00052 | M00307 | 0.59 | 8.73E-08 | 2.06E-07 |
| M00052 | M00364 | 0.51 | 7.86E-06 | 1.56E-05 |
| M00052 | M00432 | 0.60 | 3.80E-08 | 9.42E-08 |
| M00052 | M00527 | 0.80 | 2.57E-16 | 1.43E-15 |
| M00052 | M00549 | 0.70 | 3.41E-11 | 1.11E-10 |
| M00052 | M00615 | 0.29 | 0.016227 | 0.020204 |
| M00052 | M00621 | 0.77 | 1.82E-14 | 8.37E-14 |
| M00052 | M00793 | 0.70 | 2.07E-11 | 6.94E-11 |
| M00052 | M00842 | 0.71 | 1.36E-11 | 4.66E-11 |
| M00052 | M00843 | 0.72 | 3.24E-12 | 1.18E-11 |
| M00052 | M00844 | 0.71 | 1.13E-11 | 3.91E-11 |
| M00052 | M00854 | 0.26 | 0.034307 | 0.041101 |
| M00052 | M00880 | 0.66 | 1.01E-09 | 2.88E-09 |
| M00052 | M00881 | 0.79 | 1.07E-15 | 5.63E-15 |
| M00052 | M00909 | 0.81 | 2.91E-17 | 1.84E-16 |
| M00052 | M00932 | 0.52 | 5.63E-06 | 1.13E-05 |
| M00052 | M00970 | 0.55 | 9.53E-07 | 2.08E-06 |
| M00083 | M00086 | 0.44 | 0.000167 | 0.000281 |
| M00083 | M00096 | 0.67 | 3.60E-10 | 1.07E-09 |
| M00083 | M00097 | 0.90 | 3.44E-25 | 4.01E-24 |
| M00083 | M00112 | 0.75 | 1.87E-13 | 7.78E-13 |
| M00083 | M00115 | 0.52 | 3.80E-06 | 7.84E-06 |
| M00083 | M00118 | 0.49 | 2.40E-05 | 4.49E-05 |
| M00083 | M00119 | 0.94 | 6.17E-33 | 1.22E-31 |
| M00083 | M00120 | 0.70 | 2.96E-11 | 9.78E-11 |
| M00083 | M00121 | 0.96 | 1.34E-39 | 4.06E-38 |
| M00083 | M00133 | 0.80 | 3.05E-16 | 1.69E-15 |
| M00083 | M00135 | 0.92 | 1.01E-29 | 1.65E-28 |
| M00083 | M00145 | 0.59 | 9.95E-08 | 2.34E-07 |
| M00083 | M00155 | 0.91 | 7.68E-28 | 1.12E-26 |
| M00083 | M00157 | 0.43 | 0.000252 | 0.000407 |
| M00083 | M00161 | 0.33 | 0.004959 | 0.006627 |
| M00083 | M00163 | 0.35 | 0.002859 | 0.003936 |
| M00083 | M00165 | 0.68 | 1.93E-10 | 5.90E-10 |
| M00083 | M00175 | 0.33 | 0.006028 | 0.007961 |
| M00083 | M00176 | 0.85 | 4.17E-20 | 3.25E-19 |
| M00083 | M00307 | 0.46 | 6.14E-05 | 0.000109 |
| M00083 | M00364 | 0.43 | 0.000227 | 0.000371 |
| M00083 | M00432 | 0.91 | 1.45E-27 | 2.03E-26 |
| M00083 | M00527 | 0.94 | 1.32E-32 | 2.57E-31 |
| M00083 | M00549 | 0.35 | 0.003382 | 0.004613 |
| M00083 | M00621 | 0.66 | 8.10E-10 | 2.34E-09 |
| M00083 | M00793 | 0.91 | 9.77E-27 | 1.32E-25 |
| M00083 | M00842 | 0.78 | 3.98E-15 | 1.97E-14 |
| M00083 | M00843 | 0.79 | 5.79E-16 | 3.10E-15 |
| M00083 | M00844 | 0.85 | 9.42E-21 | 7.76E-20 |
| M00083 | M00854 | 0.56 | 4.74E-07 | 1.06E-06 |
| M00083 | M00880 | 0.78 | 3.38E-15 | 1.70E-14 |
| M00083 | M00881 | 0.85 | 9.41E-21 | 7.76E-20 |
| M00083 | M00909 | 0.94 | 4.81E-33 | 9.60E-32 |
| M00083 | M00932 | 0.46 | 8.52E-05 | 0.000147 |
| M00083 | M00950 | 0.53 | 3.44E-06 | 7.17E-06 |
| M00086 | M00096 | 0.52 | 3.66E-06 | 7.60E-06 |
| M00086 | M00097 | 0.52 | 5.10E-06 | 1.03E-05 |
| M00086 | M00112 | 0.68 | 1.30E-10 | 3.99E-10 |
| M00086 | M00115 | 0.60 | 3.81E-08 | 9.43E-08 |
| M00086 | M00118 | 0.49 | 1.59E-05 | 3.03E-05 |
| M00086 | M00119 | 0.43 | 0.000267 | 0.000429 |
| M00086 | M00120 | 0.52 | 5.77E-06 | 1.16E-05 |
| M00086 | M00121 | 0.54 | 1.30E-06 | 2.80E-06 |
| M00086 | M00133 | 0.52 | 5.16E-06 | 1.04E-05 |
| M00086 | M00135 | 0.30 | 0.012079 | 0.015394 |
| M00155 | M00843 | 0.81 | 3.20E-17 | 1.98E-16 |
| M00155 | M00844 | 0.78 | 1.55E-15 | 8.03E-15 |
| M00155 | M00854 | 0.30 | 0.011558 | 0.0148 |
| M00155 | M00880 | 0.79 | 4.46E-16 | 2.43E-15 |
| M00155 | M00881 | 0.90 | 3.66E-26 | 4.85E-25 |
| M00155 | M00909 | 0.90 | 7.52E-26 | 9.79E-25 |
| M00155 | M00932 | 0.38 | 0.001099 | 0.00161 |
| M00157 | M00161 | 0.33 | 0.004971 | 0.00664 |
| M00157 | M00163 | 0.46 | 6.13E-05 | 0.00011 |
| M00157 | M00165 | 0.55 | 9.56E-07 | 2.08E-06 |
| M00157 | M00175 | 0.34 | 0.004436 | 0.00596 |
| M00157 | M00176 | 0.36 | 0.002264 | 0.00316 |
| M00157 | M00307 | 0.67 | 2.66E-10 | 8.02E-10 |
| M00157 | M00432 | 0.57 | 2.64E-07 | 6.04E-07 |
| M00157 | M00527 | 0.31 | 0.010762 | 0.01387 |
| M00157 | M00549 | 0.32 | 0.007762 | 0.01011 |
| M00157 | M00621 | 0.50 | 1.04E-05 | 2.03E-05 |
| M00157 | M00842 | 0.64 | 3.37E-09 | 9.17E-09 |
| M00157 | M00843 | 0.64 | 4.41E-09 | 1.18E-08 |
| M00157 | M00844 | 0.52 | 5.07E-06 | 1.03E-05 |
| M00157 | M00854 | 0.56 | 4.50E-07 | 1.01E-06 |
| M00157 | M00880 | 0.27 | 0.023905 | 0.02909 |
| M00157 | M00881 | 0.35 | 0.002776 | 0.00383 |
| M00157 | M00909 | 0.48 | 2.58E-05 | 4.80E-05 |
| M00157 | M00950 | 0.56 | 5.92E-07 | 1.31E-06 |
| M00157 | M00970 | 0.28 | 0.018152 | 0.02243 |
| M00161 | M00165 | 0.34 | 0.004723 | 0.00633 |
| M00161 | M00175 | 0.59 | 7.62E-08 | 1.81E-07 |
| M00161 | M00176 | 0.47 | 3.96E-05 | 7.16E-05 |
| M00161 | M00307 | 0.48 | 3.36E-05 | 6.13E-05 |
| M00161 | M00364 | 0.57 | 3.78E-07 | 8.53E-07 |
| M00161 | M00432 | 0.33 | 0.005124 | 0.00683 |
| M00161 | M00527 | 0.35 | 0.002764 | 0.00382 |
| M00161 | M00549 | 0.45 | 0.00012 | 0.0002 |
| M00161 | M00621 | 0.46 | 6.53E-05 | 0.00012 |
| M00161 | M00842 | 0.33 | 0.005103 | 0.00681 |
| M00161 | M00843 | 0.36 | 0.002671 | 0.0037 |
| M00161 | M00844 | 0.44 | 0.000173 | 0.00029 |
| M00161 | M00880 | 0.25 | 0.039459 | 0.04696 |
| M00161 | M00881 | 0.50 | 1.14E-05 | 2.21E-05 |
| M00161 | M00909 | 0.39 | 0.000822 | 0.00123 |
| M00161 | M00932 | 0.50 | 1.40E-05 | 2.69E-05 |
| M00161 | M00970 | 0.37 | 0.001571 | 0.00226 |
| M00163 | M00165 | 0.86 | 6.79E-21 | 5.69E-20 |
| M00163 | M00175 | 0.31 | 0.008831 | 0.01146 |
| M00163 | M00432 | 0.66 | 5.16E-10 | 1.51E-09 |
| M00163 | M00844 | 0.48 | 3.66E-05 | 6.64E-05 |
| M00163 | M00854 | 0.93 | 5.02E-31 | 9.15E-30 |
| M00163 | M00909 | 0.36 | 0.002433 | 0.00338 |
| M00163 | M00950 | 0.96 | 7.07E-39 | 1.98E-37 |
| M00165 | M00175 | 0.46 | 7.14E-05 | 0.00013 |
| M00165 | M00176 | 0.60 | 4.67E-08 | 1.15E-07 |
| M00165 | M00307 | 0.45 | 0.000115 | 0.0002 |
| M00165 | M00364 | 0.32 | 0.006539 | 0.00862 |
| M00165 | M00432 | 0.86 | 1.16E-21 | 9.95E-21 |
| M00165 | M00527 | 0.43 | 0.000241 | 0.00039 |
| M00165 | M00549 | 0.27 | 0.025899 | 0.03139 |
| M00165 | M00621 | 0.34 | 0.004513 | 0.00606 |
| M00165 | M00793 | 0.35 | 0.003067 | 0.00421 |
| M00165 | M00842 | 0.40 | 0.000693 | 0.00105 |
| M00165 | M00843 | 0.40 | 0.00065 | 0.00099 |
| M00165 | M00844 | 0.73 | 9.45E-13 | 3.61E-12 |
| M00165 | M00854 | 0.92 | 7.67E-30 | 1.28E-28 |

| Module1 | Module2 | R | P | Padj |
|---|---|---|---|---|
| M00019 | M00176 | 0.74 | 2.64E-13 | 1.07E-12 |
| M00019 | M00307 | 0.33 | 0.005694 | 0.007561 |
| M00019 | M00364 | 0.27 | 0.02407 | 0.029269 |
| M00019 | M00432 | 0.75 | 1.41E-13 | 5.97E-13 |
| M00019 | M00527 | 0.96 | 2.07E-40 | 6.72E-39 |
| M00019 | M00549 | 0.29 | 0.016151 | 0.020127 |
| M00019 | M00621 | 0.70 | 2.80E-11 | 9.33E-11 |
| M00019 | M00793 | 1.00 | 4.06E-73 | 2.11E-71 |
| M00019 | M00842 | 0.81 | 4.42E-17 | 2.71E-16 |
| M00019 | M00843 | 0.82 | 9.78E-18 | 6.48E-17 |
| M00019 | M00844 | 0.74 | 5.03E-13 | 1.96E-12 |
| M00019 | M00854 | 0.29 | 0.015986 | 0.019955 |
| M00019 | M00880 | 0.81 | 1.80E-17 | 1.17E-16 |
| M00019 | M00881 | 0.86 | 8.01E-21 | 6.68E-20 |
| M00019 | M00909 | 0.88 | 1.87E-23 | 1.81E-22 |
| M00019 | M00932 | 0.30 | 0.012247 | 0.015568 |
| M00021 | M00023 | 0.60 | 5.22E-08 | 1.28E-07 |
| M00021 | M00024 | 0.70 | 2.96E-11 | 9.78E-11 |
| M00021 | M00028 | 0.65 | 1.05E-09 | 2.99E-09 |
| M00021 | M00048 | 0.43 | 0.000205 | 0.000338 |
| M00021 | M00049 | 0.70 | 2.01E-11 | 6.76E-11 |
| M00021 | M00050 | 0.77 | 7.26E-15 | 3.53E-14 |
| M00021 | M00052 | 0.69 | 7.60E-11 | 2.41E-10 |
| M00021 | M00083 | 0.81 | 3.55E-17 | 2.20E-16 |
| M00021 | M00086 | 0.38 | 0.001284 | 0.001871 |
| M00021 | M00096 | 0.41 | 0.000436 | 0.000687 |
| M00021 | M00097 | 0.77 | 9.68E-15 | 4.63E-14 |
| M00021 | M00112 | 0.62 | 1.55E-08 | 3.98E-08 |
| M00021 | M00115 | 0.35 | 0.003289 | 0.004499 |
| M00021 | M00118 | 0.49 | 1.83E-05 | 3.45E-05 |
| M00021 | M00119 | 0.80 | 1.25E-16 | 7.26E-16 |
| M00021 | M00120 | 0.41 | 0.000498 | 0.000775 |
| M00021 | M00121 | 0.78 | 1.49E-15 | 7.80E-15 |
| M00021 | M00133 | 0.54 | 1.62E-06 | 3.47E-06 |
| M00021 | M00135 | 0.81 | 5.37E-17 | 3.26E-16 |
| M00021 | M00145 | 0.37 | 0.002043 | 0.002868 |
| M00021 | M00155 | 0.80 | 1.37E-16 | 7.86E-16 |
| M00021 | M00157 | 0.45 | 0.000119 | 0.000203 |
| M00021 | M00161 | 0.31 | 0.010217 | 0.013182 |
| M00021 | M00165 | 0.40 | 0.000668 | 0.001009 |
| M00021 | M00176 | 0.64 | 2.37E-09 | 6.50E-09 |
| M00021 | M00307 | 0.60 | 6.92E-08 | 1.65E-07 |
| M00021 | M00364 | 0.29 | 0.014617 | 0.018356 |
| M00021 | M00432 | 0.66 | 6.66E-10 | 1.93E-09 |
| M00021 | M00527 | 0.80 | 2.00E-16 | 1.14E-15 |
| M00021 | M00549 | 0.31 | 0.009232 | 0.011954 |
| M00021 | M00621 | 0.64 | 3.56E-09 | 9.65E-09 |
| M00021 | M00793 | 0.78 | 1.69E-15 | 8.70E-15 |
| M00021 | M00842 | 0.77 | 7.17E-15 | 3.51E-14 |
| M00021 | M00843 | 0.77 | 1.06E-14 | 5.04E-14 |
| M00021 | M00844 | 0.68 | 1.20E-10 | 3.72E-10 |
| M00021 | M00854 | 0.29 | 0.01533 | 0.019219 |
| M00021 | M00880 | 0.69 | 6.91E-11 | 2.19E-10 |
| M00021 | M00881 | 0.73 | 1.13E-12 | 4.27E-12 |
| M00021 | M00909 | 0.74 | 3.16E-13 | 1.27E-12 |
| M00021 | M00932 | 0.28 | 0.021382 | 0.02611 |
| M00022 | M00023 | 0.44 | 0.000178 | 0.000297 |
| M00022 | M00024 | 0.29 | 0.015889 | 0.019851 |
| M00022 | M00028 | 0.50 | 1.09E-05 | 2.12E-05 |
| M00022 | M00048 | 0.31 | 0.009162 | 0.011874 |
| M00022 | M00049 | 0.41 | 0.000454 | 0.000711 |
| M00022 | M00050 | 0.26 | 0.032875 | 0.039506 |
| M00022 | M00052 | 0.58 | 2.12E-07 | 4.88E-07 |
| M00086 | M00145 | 0.34 | 0.004718 | 0.006328 |
| M00086 | M00155 | 0.42 | 0.000313 | 0.000501 |
| M00086 | M00157 | 0.42 | 0.000371 | 0.000587 |
| M00086 | M00161 | 0.57 | 2.52E-07 | 5.78E-07 |
| M00086 | M00165 | 0.39 | 0.000823 | 0.001227 |
| M00086 | M00175 | 0.50 | 1.09E-05 | 2.13E-05 |
| M00086 | M00176 | 0.58 | 1.67E-07 | 3.87E-07 |
| M00086 | M00307 | 0.53 | 2.63E-06 | 5.54E-06 |
| M00086 | M00364 | 0.75 | 1.80E-13 | 7.51E-13 |
| M00086 | M00432 | 0.43 | 0.000219 | 0.000359 |
| M00086 | M00527 | 0.47 | 5.04E-05 | 9.04E-05 |
| M00086 | M00549 | 0.47 | 5.10E-05 | 9.14E-05 |
| M00086 | M00621 | 0.42 | 0.000339 | 0.00054 |
| M00086 | M00793 | 0.27 | 0.024246 | 0.029459 |
| M00086 | M00842 | 0.47 | 5.14E-05 | 9.18E-05 |
| M00086 | M00843 | 0.46 | 6.05E-05 | 0.000108 |
| M00086 | M00844 | 0.50 | 1.28E-05 | 2.47E-05 |
| M00086 | M00854 | 0.30 | 0.013863 | 0.017469 |
| M00086 | M00880 | 0.29 | 0.015345 | 0.019221 |
| M00086 | M00881 | 0.49 | 1.71E-05 | 3.24E-05 |
| M00086 | M00909 | 0.46 | 7.22E-05 | 0.000127 |
| M00086 | M00932 | 0.69 | 6.23E-11 | 2.00E-10 |
| M00086 | M00950 | 0.27 | 0.024459 | 0.029668 |
| M00086 | M00970 | 0.37 | 0.001661 | 0.002374 |
| M00096 | M00097 | 0.49 | 1.76E-05 | 3.33E-05 |
| M00096 | M00112 | 0.44 | 0.000186 | 0.000308 |
| M00096 | M00115 | 0.87 | 2.24E-22 | 1.99E-21 |
| M00096 | M00118 | 0.33 | 0.005255 | 0.006998 |
| M00096 | M00119 | 0.46 | 7.95E-05 | 0.000139 |
| M00096 | M00120 | 0.94 | 1.12E-33 | 2.30E-32 |
| M00096 | M00121 | 0.63 | 7.45E-09 | 1.97E-08 |
| M00096 | M00133 | 0.91 | 9.09E-27 | 1.24E-25 |
| M00096 | M00135 | 0.41 | 0.000507 | 0.000787 |
| M00096 | M00145 | 0.76 | 4.58E-14 | 2.00E-13 |
| M00096 | M00155 | 0.44 | 0.000179 | 0.000298 |
| M00096 | M00157 | 0.52 | 5.08E-06 | 1.03E-05 |
| M00096 | M00161 | 0.40 | 0.000598 | 0.000914 |
| M00096 | M00163 | 0.77 | 9.55E-15 | 4.58E-14 |
| M00096 | M00165 | 0.88 | 8.45E-24 | 8.62E-23 |
| M00096 | M00175 | 0.46 | 6.69E-05 | 0.000118 |
| M00096 | M00176 | 0.60 | 6.55E-08 | 1.58E-07 |
| M00096 | M00307 | 0.45 | 0.00011 | 0.000188 |
| M00096 | M00364 | 0.48 | 2.81E-05 | 5.21E-05 |
| M00096 | M00432 | 0.85 | 4.40E-20 | 3.41E-19 |
| M00096 | M00527 | 0.48 | 3.63E-05 | 6.60E-05 |
| M00096 | M00549 | 0.26 | 0.028428 | 0.034396 |
| M00096 | M00621 | 0.35 | 0.003343 | 0.004568 |
| M00096 | M00793 | 0.36 | 0.002221 | 0.003105 |
| M00096 | M00842 | 0.51 | 6.60E-06 | 1.32E-05 |
| M00096 | M00843 | 0.51 | 8.07E-06 | 1.59E-05 |
| M00096 | M00844 | 0.77 | 1.76E-14 | 8.09E-14 |
| M00096 | M00854 | 0.85 | 2.70E-20 | 2.14E-19 |
| M00096 | M00880 | 0.37 | 0.001859 | 0.002634 |
| M00096 | M00881 | 0.51 | 7.55E-06 | 1.50E-05 |
| M00096 | M00909 | 0.65 | 1.57E-09 | 4.35E-09 |
| M00096 | M00932 | 0.40 | 0.000768 | 0.001151 |
| M00096 | M00950 | 0.84 | 1.54E-19 | 1.16E-18 |
| M00097 | M00112 | 0.77 | 6.41E-15 | 3.15E-14 |
| M00097 | M00115 | 0.45 | 0.000121 | 0.000205 |
| M00097 | M00118 | 0.54 | 1.62E-06 | 3.48E-06 |
| M00097 | M00119 | 0.91 | 5.35E-28 | 8.13E-27 |
| M00097 | M00120 | 0.50 | 1.48E-05 | 2.84E-05 |
| M00097 | M00121 | 0.88 | 1.07E-23 | 1.08E-22 |
| M00165 | M00880 | 0.37 | 0.001762 | 0.00251 |
| M00165 | M00881 | 0.48 | 2.48E-05 | 4.62E-05 |
| M00165 | M00909 | 0.67 | 3.70E-10 | 1.10E-09 |
| M00165 | M00932 | 0.41 | 0.000451 | 0.00071 |
| M00165 | M00950 | 0.94 | 4.27E-33 | 8.65E-32 |
| M00175 | M00176 | 0.32 | 0.006548 | 0.00862 |
| M00175 | M00307 | 0.30 | 0.012309 | 0.01563 |
| M00175 | M00364 | 0.39 | 0.001032 | 0.00152 |
| M00175 | M00432 | 0.43 | 0.000204 | 0.00034 |
| M00175 | M00527 | 0.26 | 0.032786 | 0.03944 |
| M00175 | M00549 | 0.74 | 4.48E-13 | 1.76E-12 |
| M00175 | M00615 | 0.50 | 9.95E-06 | 1.95E-05 |
| M00175 | M00621 | 0.42 | 0.000343 | 0.00055 |
| M00175 | M00844 | 0.51 | 8.23E-06 | 1.62E-05 |
| M00175 | M00854 | 0.49 | 2.22E-05 | 4.16E-05 |
| M00175 | M00881 | 0.40 | 0.000779 | 0.00117 |
| M00175 | M00909 | 0.50 | 9.69E-06 | 1.90E-05 |
| M00175 | M00932 | 0.30 | 0.01277 | 0.01618 |
| M00175 | M00950 | 0.43 | 0.000272 | 0.00044 |
| M00175 | M00970 | 0.67 | 3.10E-10 | 9.30E-10 |
| M00176 | M00307 | 0.52 | 5.73E-06 | 1.15E-05 |
| M00176 | M00364 | 0.73 | 1.83E-12 | 6.81E-12 |
| M00176 | M00432 | 0.75 | 1.53E-13 | 6.42E-13 |
| M00176 | M00527 | 0.85 | 1.19E-20 | 9.72E-20 |
| M00176 | M00549 | 0.33 | 0.005865 | 0.00776 |
| M00176 | M00621 | 0.57 | 2.45E-07 | 5.63E-07 |
| M00176 | M00793 | 0.72 | 2.05E-12 | 7.59E-12 |
| M00176 | M00842 | 0.63 | 7.80E-09 | 2.05E-08 |
| M00176 | M00843 | 0.66 | 5.36E-10 | 1.56E-09 |
| M00176 | M00844 | 0.72 | 3.90E-12 | 1.40E-11 |
| M00176 | M00854 | 0.40 | 0.000777 | 0.00116 |
| M00176 | M00880 | 0.60 | 5.41E-08 | 1.32E-07 |
| M00176 | M00881 | 0.80 | 1.32E-16 | 7.61E-16 |
| M00176 | M00909 | 0.77 | 1.19E-14 | 5.63E-14 |
| M00176 | M00932 | 0.78 | 1.50E-15 | 7.82E-15 |
| M00176 | M00950 | 0.41 | 0.000434 | 0.00068 |
| M00307 | M00364 | 0.40 | 0.000738 | 0.00111 |
| M00307 | M00432 | 0.48 | 3.15E-05 | 5.78E-05 |
| M00307 | M00527 | 0.41 | 0.000507 | 0.00079 |
| M00307 | M00549 | 0.37 | 0.001998 | 0.00281 |
| M00307 | M00621 | 0.54 | 1.82E-06 | 3.88E-06 |
| M00307 | M00793 | 0.28 | 0.02011 | 0.02464 |
| M00307 | M00842 | 0.52 | 3.64E-06 | 7.57E-06 |
| M00307 | M00843 | 0.54 | 1.96E-06 | 4.17E-06 |
| M00307 | M00844 | 0.49 | 1.89E-05 | 3.55E-05 |
| M00307 | M00854 | 0.37 | 0.001573 | 0.00226 |
| M00307 | M00880 | 0.34 | 0.003837 | 0.00518 |
| M00307 | M00881 | 0.46 | 8.47E-05 | 0.00015 |
| M00307 | M00909 | 0.47 | 4.67E-05 | 8.40E-05 |
| M00307 | M00932 | 0.37 | 0.001651 | 0.00236 |
| M00307 | M00950 | 0.37 | 0.001703 | 0.00243 |
| M00307 | M00970 | 0.30 | 0.01195 | 0.01526 |
| M00364 | M00432 | 0.35 | 0.003271 | 0.00448 |
| M00364 | M00527 | 0.50 | 9.77E-06 | 1.92E-05 |
| M00364 | M00549 | 0.35 | 0.003232 | 0.00443 |
| M00364 | M00621 | 0.32 | 0.0071 | 0.0093 |
| M00364 | M00793 | 0.26 | 0.031413 | 0.03785 |
| M00364 | M00842 | 0.30 | 0.011836 | 0.01512 |
| M00364 | M00843 | 0.33 | 0.005757 | 0.00763 |
| M00364 | M00844 | 0.39 | 0.000972 | 0.00144 |
| M00364 | M00881 | 0.51 | 7.89E-06 | 1.56E-05 |
| M00364 | M00909 | 0.37 | 0.001633 | 0.00234 |
| M00364 | M00932 | 0.89 | 3.53E-25 | 4.09E-24 |

| Module1 | Module2 | R | P | Padj | Module1 | Module2 | R | P | Padj | Module1 | Module2 | R | P | Padj |
|---|---|---|---|---|---|---|---|---|---|---|---|---|---|---|
| M00022 | M00086 | 0.41 | 0.00054 | 0.000832 | M00097 | M00133 | 0.66 | 9.68E-10 | 2.77E-09 | M00432 | M00527 | 0.75 | 1.89E-13 | 7.81E-13 |
| M00022 | M00097 | 0.52 | 3.87E-06 | 7.97E-06 | M00097 | M00135 | 0.90 | 2.49E-25 | 2.98E-24 | M00432 | M00549 | 0.36 | 0.002606 | 0.00362 |
| M00022 | M00112 | 0.27 | 0.022758 | 0.02772 | M00097 | M00145 | 0.50 | 1.44E-05 | 2.76E-05 | M00432 | M00621 | 0.55 | 8.79E-07 | 1.93E-06 |
| M00022 | M00115 | 0.34 | 0.004205 | 0.005666 | M00097 | M00155 | 0.93 | 2.80E-30 | 4.80E-29 | M00432 | M00793 | 0.71 | 1.23E-11 | 4.23E-11 |
| M00022 | M00118 | 0.41 | 0.000541 | 0.000832 | M00097 | M00157 | 0.45 | 0.000113 | 0.000192 | M00432 | M00842 | 0.68 | 1.60E-10 | 4.91E-10 |
| M00022 | M00133 | 0.28 | 0.020851 | 0.025482 | M00097 | M00161 | 0.46 | 7.97E-05 | 0.000139 | M00432 | M00843 | 0.68 | 1.21E-10 | 3.73E-10 |
| M00022 | M00145 | 0.37 | 0.001882 | 0.002663 | M00097 | M00165 | 0.48 | 3.01E-05 | 5.57E-05 | M00432 | M00844 | 0.88 | 3.24E-23 | 3.07E-22 |
| M00022 | M00155 | 0.26 | 0.03112 | 0.03753 | M00097 | M00175 | 0.49 | 1.88E-05 | 3.54E-05 | M00432 | M00854 | 0.82 | 1.56E-17 | 1.02E-16 |
| M00022 | M00157 | 0.31 | 0.009864 | 0.012762 | M00097 | M00176 | 0.78 | 3.75E-15 | 1.87E-14 | M00432 | M00880 | 0.62 | 1.03E-08 | 2.68E-08 |
| M00022 | M00161 | 0.45 | 0.000104 | 0.000178 | M00097 | M00307 | 0.49 | 1.71E-05 | 3.24E-05 | M00432 | M00881 | 0.71 | 7.34E-12 | 2.57E-11 |
| M00022 | M00175 | 0.75 | 2.05E-13 | 8.42E-13 | M00097 | M00364 | 0.45 | 0.000101 | 0.000173 | M00432 | M00909 | 0.90 | 2.72E-26 | 3.63E-25 |
| M00022 | M00307 | 0.32 | 0.00695 | 0.009121 | M00097 | M00432 | 0.77 | 9.08E-15 | 4.37E-14 | M00432 | M00932 | 0.36 | 0.002635 | 0.00365 |
| M00022 | M00364 | 0.29 | 0.016411 | 0.020415 | M00097 | M00527 | 0.92 | 9.26E-30 | 1.53E-28 | M00432 | M00950 | 0.80 | 2.25E-16 | 1.27E-15 |
| M00022 | M00549 | 0.95 | 2.08E-35 | 4.88E-34 | M00097 | M00549 | 0.63 | 7.25E-09 | 1.92E-08 | M00527 | M00549 | 0.36 | 0.002421 | 0.00337 |
| M00022 | M00615 | 0.60 | 6.15E-08 | 1.48E-07 | M00097 | M00615 | 0.28 | 0.020319 | 0.024854 | M00527 | M00621 | 0.72 | 3.75E-12 | 1.35E-11 |
| M00022 | M00621 | 0.48 | 3.30E-05 | 6.05E-05 | M00097 | M00621 | 0.81 | 3.00E-17 | 1.88E-16 | M00527 | M00793 | 0.96 | 9.96E-39 | 2.74E-37 |
| M00022 | M00844 | 0.39 | 0.000914 | 0.001356 | M00097 | M00793 | 0.87 | 2.25E-22 | 1.99E-21 | M00527 | M00842 | 0.81 | 1.88E-17 | 1.22E-16 |
| M00022 | M00854 | 0.26 | 0.032527 | 0.039161 | M00097 | M00842 | 0.80 | 2.93E-16 | 1.63E-15 | M00527 | M00843 | 0.83 | 7.26E-19 | 5.24E-18 |
| M00022 | M00881 | 0.32 | 0.006695 | 0.008794 | M00097 | M00843 | 0.81 | 2.91E-17 | 1.84E-16 | M00527 | M00844 | 0.75 | 1.50E-13 | 6.32E-13 |
| M00022 | M00909 | 0.48 | 3.56E-05 | 6.47E-05 | M00097 | M00844 | 0.81 | 6.52E-17 | 3.91E-16 | M00527 | M00854 | 0.28 | 0.018461 | 0.02275 |
| M00022 | M00970 | 0.95 | 7.40E-36 | 1.86E-34 | M00097 | M00854 | 0.39 | 0.000977 | 0.001446 | M00527 | M00880 | 0.79 | 5.66E-16 | 3.04E-15 |
| M00023 | M00024 | 0.70 | 2.90E-11 | 9.62E-11 | M00097 | M00880 | 0.75 | 6.77E-14 | 2.91E-13 | M00527 | M00881 | 0.90 | 1.55E-25 | 1.95E-24 |
| M00023 | M00028 | 0.73 | 1.41E-12 | 5.27E-12 | M00097 | M00881 | 0.88 | 1.29E-23 | 1.29E-22 | M00527 | M00909 | 0.88 | 2.08E-23 | 1.99E-22 |
| M00023 | M00048 | 0.68 | 8.88E-11 | 2.80E-10 | M00097 | M00909 | 0.95 | 1.13E-35 | 2.75E-34 | M00527 | M00932 | 0.50 | 1.09E-05 | 2.12E-05 |
| M00023 | M00049 | 0.66 | 6.40E-10 | 1.86E-09 | M00097 | M00932 | 0.43 | 0.000269 | 0.000432 | M00549 | M00615 | 0.60 | 6.14E-08 | 1.48E-07 |
| M00023 | M00050 | 0.73 | 9.51E-13 | 3.62E-12 | M00097 | M00950 | 0.31 | 0.008713 | 0.011323 | M00549 | M00621 | 0.55 | 1.27E-06 | 2.74E-06 |
| M00023 | M00052 | 0.76 | 3.81E-14 | 1.71E-13 | M00097 | M00970 | 0.43 | 0.000215 | 0.000352 | M00549 | M00793 | 0.25 | 0.037568 | 0.04482 |
| M00023 | M00083 | 0.65 | 1.37E-09 | 3.82E-09 | M00112 | M00115 | 0.50 | 1.43E-05 | 2.75E-05 | M00549 | M00842 | 0.29 | 0.017574 | 0.02175 |
| M00023 | M00086 | 0.64 | 3.69E-09 | 9.94E-09 | M00112 | M00118 | 0.70 | 1.83E-11 | 6.18E-11 | M00549 | M00843 | 0.28 | 0.018777 | 0.02312 |
| M00023 | M00096 | 0.60 | 6.74E-08 | 1.61E-07 | M00112 | M00119 | 0.78 | 1.93E-15 | 9.89E-15 | M00549 | M00844 | 0.52 | 3.98E-06 | 8.16E-06 |
| M00023 | M00097 | 0.72 | 2.73E-12 | 1.00E-11 | M00112 | M00120 | 0.50 | 1.15E-05 | 2.24E-05 | M00549 | M00854 | 0.34 | 0.004833 | 0.00647 |
| M00023 | M00112 | 0.74 | 4.17E-13 | 1.65E-12 | M00112 | M00121 | 0.85 | 2.13E-20 | 1.72E-19 | M00549 | M00880 | 0.42 | 0.000353 | 0.00056 |
| M00023 | M00115 | 0.64 | 2.94E-09 | 8.01E-09 | M00112 | M00133 | 0.55 | 8.98E-07 | 1.97E-06 | M00549 | M00881 | 0.46 | 7.77E-05 | 0.00014 |
| M00023 | M00118 | 0.69 | 6.68E-11 | 2.13E-10 | M00112 | M00135 | 0.70 | 1.41E-11 | 4.80E-11 | M00549 | M00909 | 0.60 | 5.62E-08 | 1.37E-07 |
| M00023 | M00119 | 0.63 | 7.45E-09 | 1.97E-08 | M00112 | M00145 | 0.34 | 0.00425 | 0.005722 | M00549 | M00932 | 0.26 | 0.033824 | 0.04056 |
| M00023 | M00120 | 0.59 | 1.20E-07 | 2.82E-07 | M00112 | M00155 | 0.74 | 2.81E-13 | 1.14E-12 | M00549 | M00970 | 0.90 | 2.29E-25 | 2.76E-24 |
| M00023 | M00121 | 0.70 | 2.98E-11 | 9.82E-11 | M00112 | M00157 | 0.36 | 0.002381 | 0.003316 | M00615 | M00970 | 0.60 | 5.40E-08 | 1.32E-07 |
| M00023 | M00133 | 0.70 | 1.54E-11 | 5.25E-11 | M00112 | M00161 | 0.52 | 3.69E-06 | 7.64E-06 | M00621 | M00793 | 0.67 | 3.07E-10 | 9.24E-10 |
| M00023 | M00135 | 0.60 | 4.19E-08 | 1.03E-07 | M00112 | M00165 | 0.40 | 0.000651 | 0.000987 | M00621 | M00842 | 0.72 | 5.08E-12 | 1.81E-11 |
| M00023 | M00145 | 0.68 | 1.25E-10 | 3.85E-10 | M00112 | M00175 | 0.36 | 0.002074 | 0.002904 | M00621 | M00843 | 0.72 | 2.66E-12 | 9.76E-12 |
| M00023 | M00155 | 0.66 | 8.51E-10 | 2.44E-09 | M00112 | M00176 | 0.85 | 1.05E-20 | 8.58E-20 | M00621 | M00844 | 0.61 | 3.16E-08 | 7.84E-08 |
| M00023 | M00157 | 0.51 | 8.07E-06 | 1.59E-05 | M00112 | M00307 | 0.51 | 7.38E-06 | 1.47E-05 | M00621 | M00854 | 0.30 | 0.01128 | 0.01448 |
| M00023 | M00161 | 0.58 | 2.02E-07 | 4.69E-07 | M00112 | M00364 | 0.71 | 6.53E-12 | 2.29E-11 | M00621 | M00880 | 0.58 | 1.34E-07 | 3.13E-07 |
| M00023 | M00165 | 0.59 | 7.06E-08 | 1.68E-07 | M00112 | M00432 | 0.58 | 2.07E-07 | 4.79E-07 | M00621 | M00881 | 0.88 | 8.58E-23 | 7.87E-22 |
| M00023 | M00175 | 0.47 | 4.90E-05 | 8.80E-05 | M00112 | M00527 | 0.82 | 7.59E-18 | 5.13E-17 | M00621 | M00909 | 0.73 | 8.33E-13 | 3.20E-12 |
| M00023 | M00176 | 0.71 | 6.42E-12 | 2.26E-11 | M00112 | M00549 | 0.45 | 9.53E-05 | 0.000164 | M00621 | M00932 | 0.25 | 0.038083 | 0.0454 |
| M00023 | M00307 | 0.61 | 3.05E-08 | 7.61E-08 | M00112 | M00621 | 0.63 | 6.53E-09 | 1.74E-08 | M00621 | M00970 | 0.43 | 0.000244 | 0.0004 |
| M00023 | M00364 | 0.51 | 6.71E-06 | 1.34E-05 | M00112 | M00793 | 0.68 | 1.17E-10 | 3.63E-10 | M00793 | M00842 | 0.79 | 5.52E-16 | 2.98E-15 |
| M00023 | M00432 | 0.63 | 6.81E-09 | 1.81E-08 | M00112 | M00842 | 0.64 | 4.13E-09 | 1.11E-08 | M00793 | M00843 | 0.80 | 1.31E-16 | 7.58E-16 |
| M00023 | M00527 | 0.65 | 1.98E-09 | 5.46E-09 | M00112 | M00843 | 0.67 | 2.13E-10 | 6.48E-10 | M00793 | M00844 | 0.70 | 2.61E-11 | 8.70E-11 |
| M00023 | M00549 | 0.56 | 7.02E-07 | 1.54E-06 | M00112 | M00844 | 0.66 | 4.88E-10 | 1.44E-09 | M00793 | M00880 | 0.81 | 4.94E-17 | 3.02E-16 |
| M00023 | M00621 | 0.67 | 3.92E-10 | 1.16E-09 | M00112 | M00854 | 0.26 | 0.029897 | 0.036115 | M00793 | M00881 | 0.84 | 3.01E-19 | 2.23E-18 |
| M00023 | M00793 | 0.53 | 3.08E-06 | 6.43E-06 | M00112 | M00880 | 0.77 | 1.62E-14 | 7.46E-14 | M00793 | M00909 | 0.85 | 3.57E-20 | 2.80E-19 |
| M00023 | M00842 | 0.67 | 2.31E-10 | 6.99E-10 | M00112 | M00881 | 0.78 | 3.43E-15 | 1.72E-14 | M00793 | M00932 | 0.29 | 0.015636 | 0.01955 |
| M00023 | M00843 | 0.69 | 6.34E-11 | 2.03E-10 | M00112 | M00909 | 0.70 | 2.88E-11 | 9.56E-11 | M00842 | M00843 | 0.99 | 2.24E-65 | 1.13E-63 |
| M00023 | M00844 | 0.71 | 9.80E-12 | 3.41E-11 | M00112 | M00932 | 0.74 | 3.99E-13 | 1.59E-12 | M00842 | M00844 | 0.69 | 4.49E-11 | 1.46E-10 |
| M00023 | M00854 | 0.44 | 0.000178 | 0.000298 | M00115 | M00118 | 0.43 | 0.000237 | 0.000385 | M00842 | M00854 | 0.29 | 0.013985 | 0.01761 |
| M00023 | M00880 | 0.61 | 2.22E-08 | 5.62E-08 | M00115 | M00119 | 0.32 | 0.008219 | 0.01069 | M00842 | M00880 | 0.67 | 3.52E-10 | 1.05E-09 |
| M00023 | M00881 | 0.72 | 1.96E-12 | 7.26E-12 | M00115 | M00120 | 0.87 | 6.66E-22 | 5.78E-21 | M00842 | M00881 | 0.75 | 2.05E-13 | 8.42E-13 |
| M00023 | M00909 | 0.70 | 2.01E-11 | 6.76E-11 | M00115 | M00121 | 0.54 | 1.86E-06 | 3.94E-06 | M00842 | M00909 | 0.76 | 6.49E-14 | 2.80E-13 |
| M00023 | M00932 | 0.56 | 4.99E-07 | 1.11E-06 | M00115 | M00133 | 0.83 | 1.06E-18 | 7.64E-18 | M00842 | M00932 | 0.25 | 0.040168 | 0.04769 |
| M00023 | M00950 | 0.37 | 0.001791 | 0.002546 | M00115 | M00135 | 0.26 | 0.032895 | 0.039506 | M00843 | M00844 | 0.69 | 6.56E-11 | 2.09E-10 |
| M00023 | M00970 | 0.41 | 0.000487 | 0.00076 | M00115 | M00145 | 0.67 | 3.43E-10 | 1.02E-09 | M00843 | M00854 | 0.29 | 0.016687 | 0.02071 |

| Module1 | Module2 | R | P | Padj |
|---|---|---|---|---|
| M00024 | M00028 | 0.60 | 3.84E-08 | 9.50E-08 |
| M00024 | M00048 | 0.46 | 7.07E-05 | 0.000124 |
| M00024 | M00049 | 0.57 | 2.76E-07 | 6.28E-07 |
| M00024 | M00050 | 0.71 | 1.17E-11 | 4.01E-11 |
| M00024 | M00052 | 0.62 | 9.89E-09 | 2.58E-08 |
| M00024 | M00083 | 0.82 | 5.26E-18 | 3.64E-17 |
| M00024 | M00086 | 0.42 | 0.000337 | 0.000538 |
| M00024 | M00096 | 0.76 | 3.95E-14 | 1.76E-13 |
| M00024 | M00097 | 0.74 | 3.01E-13 | 1.21E-12 |
| M00024 | M00112 | 0.52 | 3.97E-06 | 8.15E-06 |
| M00024 | M00115 | 0.73 | 1.26E-12 | 4.75E-12 |
| M00024 | M00118 | 0.41 | 0.000511 | 0.000793 |
| M00024 | M00119 | 0.68 | 9.33E-11 | 2.93E-10 |
| M00024 | M00120 | 0.77 | 1.15E-14 | 5.42E-14 |
| M00024 | M00121 | 0.74 | 2.42E-13 | 9.90E-13 |
| M00024 | M00133 | 0.90 | 2.55E-25 | 3.02E-24 |
| M00024 | M00135 | 0.69 | 3.74E-11 | 1.21E-10 |
| M00024 | M00145 | 0.72 | 1.92E-12 | 7.15E-12 |
| M00024 | M00155 | 0.70 | 1.90E-11 | 6.40E-11 |
| M00024 | M00157 | 0.65 | 1.78E-09 | 4.90E-09 |
| M00024 | M00161 | 0.33 | 0.004915 | 0.006574 |
| M00024 | M00163 | 0.57 | 2.88E-07 | 6.55E-07 |

| Module1 | Module2 | R | P | Padj |
|---|---|---|---|---|
| M00115 | M00155 | 0.33 | 0.006053 | 0.007987 |
| M00115 | M00157 | 0.65 | 1.99E-09 | 5.48E-09 |
| M00115 | M00161 | 0.45 | 9.15E-05 | 0.000158 |
| M00115 | M00163 | 0.68 | 1.17E-10 | 3.63E-10 |
| M00115 | M00165 | 0.80 | 3.07E-16 | 1.70E-15 |
| M00115 | M00175 | 0.55 | 1.14E-06 | 2.47E-06 |
| M00115 | M00176 | 0.58 | 1.64E-07 | 3.81E-07 |
| M00115 | M00307 | 0.61 | 2.38E-08 | 5.98E-08 |
| M00115 | M00364 | 0.53 | 2.55E-06 | 5.39E-06 |
| M00115 | M00432 | 0.71 | 6.09E-12 | 2.14E-11 |
| M00115 | M00527 | 0.36 | 0.002707 | 0.003741 |
| M00115 | M00549 | 0.41 | 0.00046 | 0.00072 |
| M00115 | M00621 | 0.40 | 0.000572 | 0.000877 |
| M00115 | M00842 | 0.42 | 0.000338 | 0.000538 |
| M00115 | M00843 | 0.43 | 0.000203 | 0.000336 |
| M00115 | M00844 | 0.69 | 5.43E-11 | 1.75E-10 |
| M00115 | M00854 | 0.79 | 8.30E-16 | 4.39E-15 |
| M00115 | M00880 | 0.29 | 0.014533 | 0.018266 |
| M00115 | M00881 | 0.45 | 9.40E-05 | 0.000162 |
| M00115 | M00909 | 0.55 | 9.58E-07 | 2.09E-06 |
| M00115 | M00932 | 0.45 | 0.0001 | 0.000172 |
| M00115 | M00950 | 0.77 | 1.20E-14 | 5.64E-14 |

| Module1 | Module2 | R | P | Padj |
|---|---|---|---|---|
| M00843 | M00880 | 0.68 | 8.65E-11 | 2.74E-10 |
| M00843 | M00881 | 0.76 | 4.35E-14 | 1.93E-13 |
| M00843 | M00909 | 0.76 | 3.79E-14 | 1.71E-13 |
| M00843 | M00932 | 0.28 | 0.018849 | 0.02319 |
| M00844 | M00854 | 0.69 | 6.43E-11 | 2.06E-10 |
| M00844 | M00880 | 0.72 | 3.53E-12 | 1.28E-11 |
| M00844 | M00881 | 0.74 | 4.76E-13 | 1.86E-12 |
| M00844 | M00909 | 0.90 | 5.97E-26 | 7.85E-25 |
| M00844 | M00932 | 0.37 | 0.001905 | 0.00269 |
| M00844 | M00950 | 0.61 | 2.04E-08 | 5.16E-08 |
| M00844 | M00970 | 0.30 | 0.011201 | 0.01439 |
| M00854 | M00880 | 0.34 | 0.00409 | 0.00552 |
| M00854 | M00881 | 0.38 | 0.001412 | 0.00205 |
| M00854 | M00909 | 0.61 | 2.88E-08 | 7.21E-08 |
| M00854 | M00950 | 0.96 | 4.16E-40 | 1.32E-38 |
| M00880 | M00881 | 0.70 | 1.43E-11 | 4.88E-11 |
| M00880 | M00909 | 0.76 | 2.44E-14 | 1.11E-13 |
| M00881 | M00909 | 0.84 | 7.19E-20 | 5.51E-19 |
| M00881 | M00932 | 0.48 | 3.04E-05 | 5.61E-05 |
| M00881 | M00950 | 0.28 | 0.019513 | 0.02395 |
| M00909 | M00932 | 0.37 | 0.001847 | 0.00262 |
| M00909 | M00950 | 0.54 | 2.10E-06 | 4.44E-06 |
| M00909 | M00970 | 0.38 | 0.001128 | 0.00165 |

### Supplementary Figures

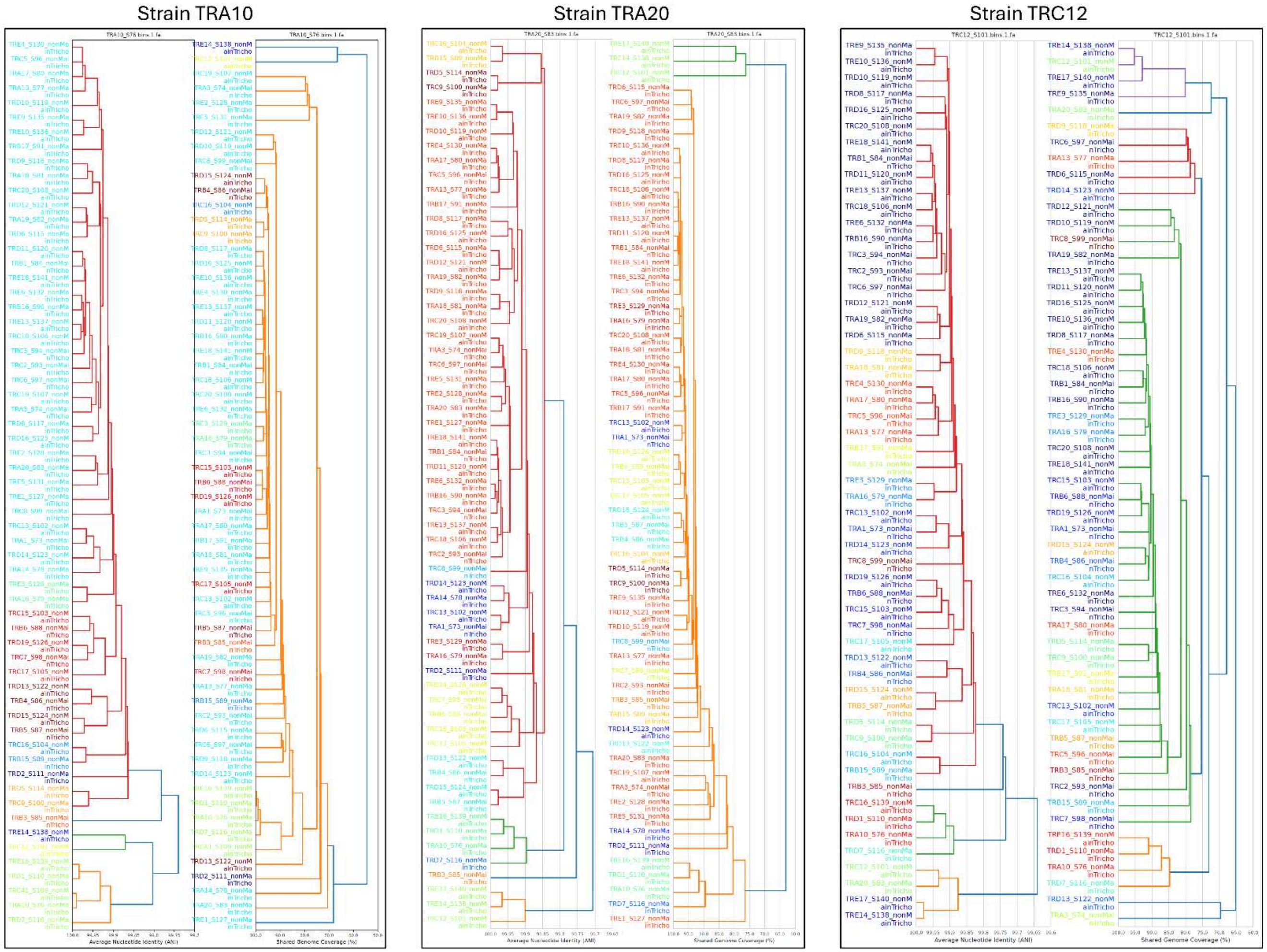


**Figure S1.** Strain diversity analysis conducted with InStrain[1] on the three (dereplicated) Trichodesmium strains.

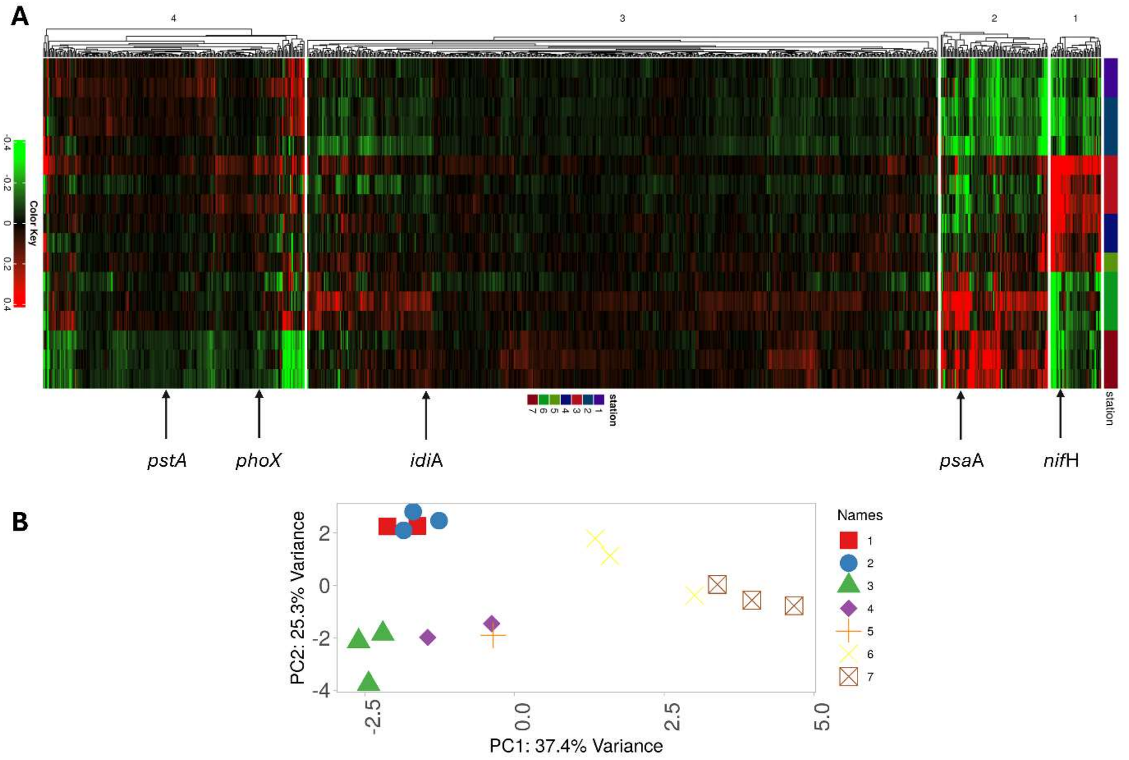


**Figure S2.** Clustering (A) and PCA (B) analyses of the TPM data from Cerdan-Garcia et al[2] as provided by the authors and normalized to the *rpo*B values. The figures above are comparable to the results presented by the authors in Figure 3 of their paper. The clustering is of the 1000 most variable genes. The *isi*A and *isi*B genes highlighted by the original authors in their Figure 3, were not included in the top 1000 genes. The figures were generated using iDEP 2 (http://bioinformatics.sdstate.edu/idep/) [3]**.**

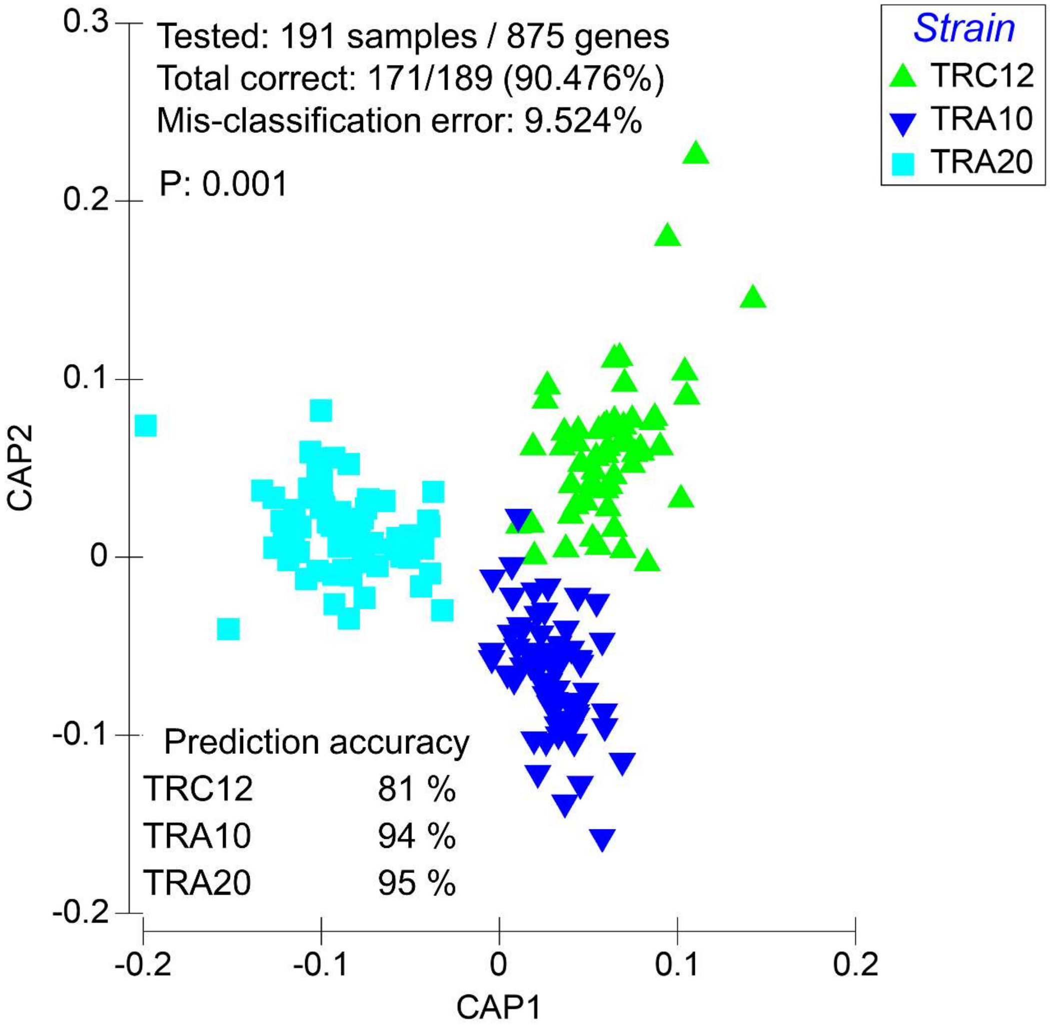


**Figure S3.** Canonical Correspondence Analysis of 875 identifiable genes across the three strains of *Trichodesmium*. The prediction accuracy remains high also when a larger repertoire of genes is used with the three strains having a clear transcriptional profile (p=0.001; 999 permutations)/

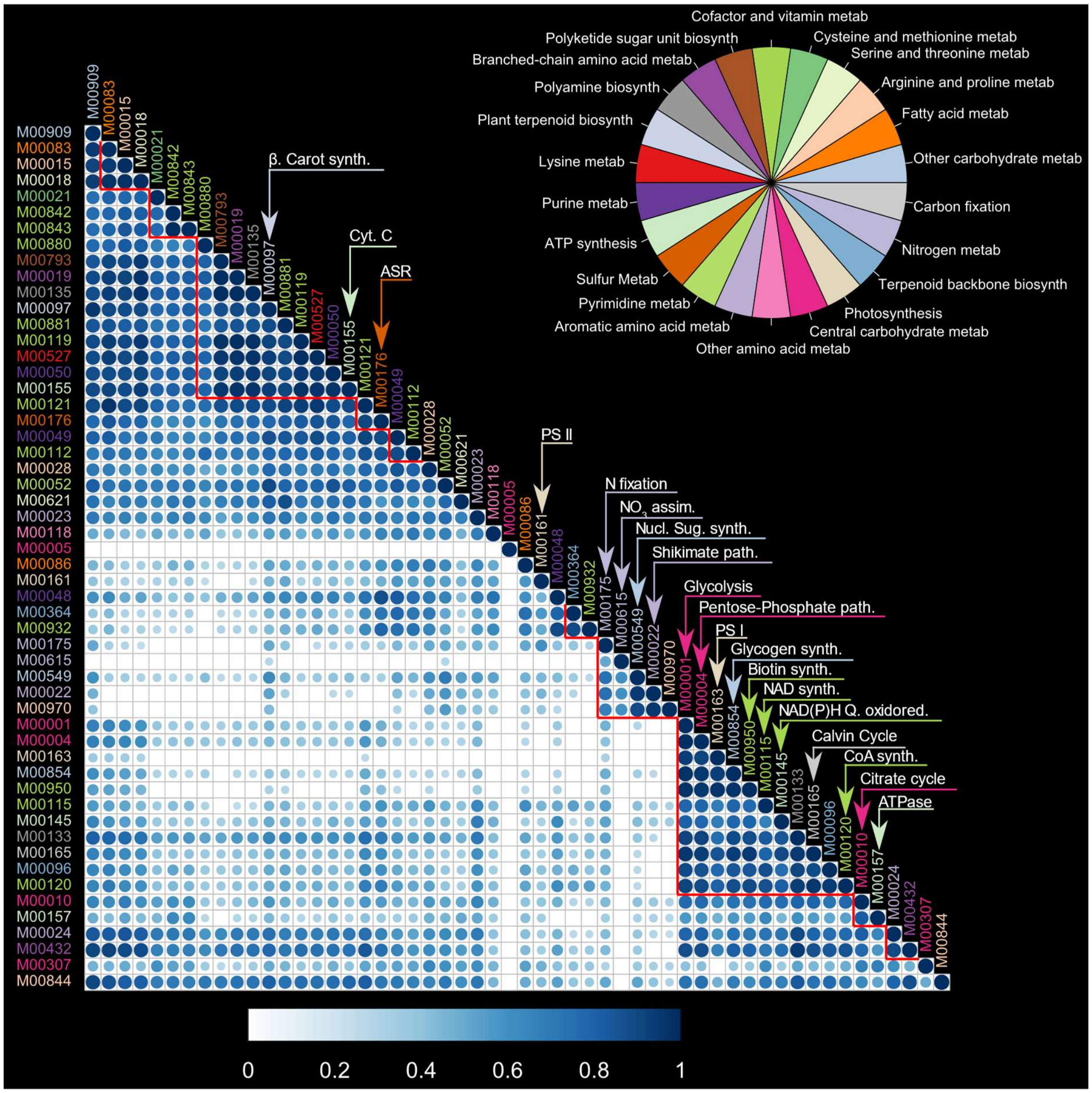


**Figure S4.** Correlation matrix of KEGG modules expression profiles. Module names are colored according to the KEGG pathway category of which the module is part of, as depicted in the color legend. The wheel of the color legend bears no quantitative information. Only significant correlations are depicted (Benjamini Hochberg false discovery rate <0.05). The Pearson correlation coefficient (R) is represented by the size, and color intensity of the blue circles. Modules whose correlation patterns form clusters are marked with red lines ($p<0.05$; number of bootstraps 10,000). For space considerations, only a selected number of modules are named in the figure. The complete list of modules correlation data is shown in Table S4, respectively. Abbreviations used in the figure: metab: metabolism; (bio)synth: (bio)synthesis.

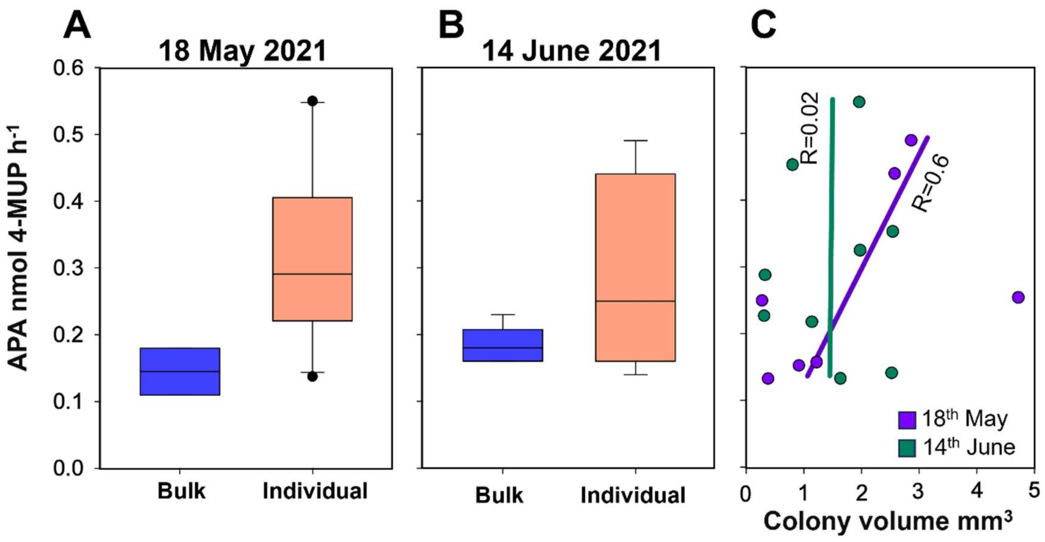


**Figure S5.** Alkaline phosphatase activity measurements for bulk samples and individual colonies in May 2021 (A,B) depict opposite misrepresentations of bulk analyses as compared to multiple individual colonies. The activity for the colony samples on the 18th of May and 14th of June is shown in relation to colony volume (C). A significant correlation between activity and size is seen only on one date.

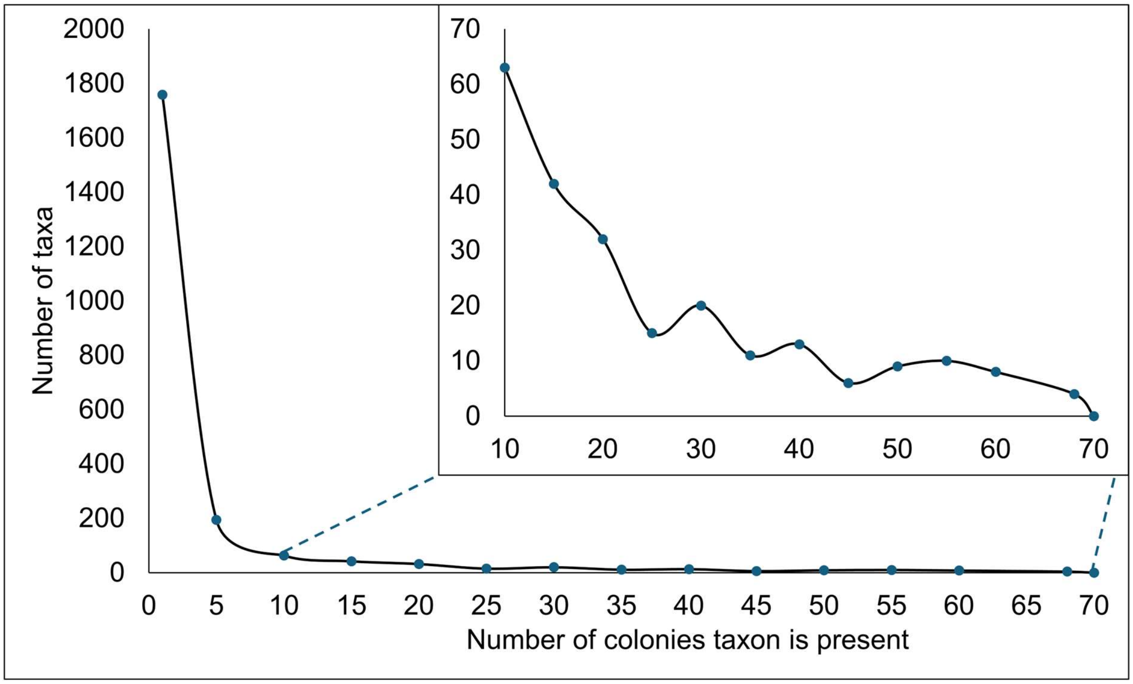


**Figure S6** Histogram of bacterial and archaeal taxa presence across the 69 *Trichodesmium* colonies, excluding Cyanobacteria. The insert focuses on taxa present in more than 10 colonies.
